# Quantifying the contribution of DNA conformational flexibility to transcription factor binding on nucleosomal DNA uncovers indirect readout across diverse TF families

**DOI:** 10.1101/2025.05.21.655105

**Authors:** Upalabdha Dey, Gustavo Sganzerla Martinez, Ravi Kumar, Venkata Rajesh Yella, Aditya Kumar

## Abstract

**Background:** Eukaryotic gene regulation depends on transcription factors (TFs) recognizing short DNA motifs within chromatin. Many of these motifs lie within nucleosomes, where DNA is sharply bent, rotationally phased, and constrained by histone-DNA contacts. Yet only a subset is occupied in any cellular context. Motif identity alone, therefore, cannot fully explain selective TF engagement with nucleosomal DNA. We asked whether sequence-derived DNA conformational flexibility provides an interpretable representation of sequence context relevant to TF recognition on nucleosomes.

**Results:** We compiled five DNA flexibility descriptors in the Python package DNAflexpy, representing bendability, torsional deformation, backbone conformational variability, and stiffness. We built quantitative models of TF binding affinity across 226 datasets from a high-throughput in vitro TF-nucleosome binding assay. Flexibility-augmented models improved prediction over mononucleotide baselines in most datasets, with smaller but reproducible gains over trinucleotide baselines. The gains were not uniform: they varied across TF families and were concordant with DNA shape-fluctuation features, suggesting that DNAflexpy descriptors capture a sequence-encoded structural signal. In PIONEAR-seq data, model performance generalized across nucleosomal templates in a TF- and sequence-dependent manner. Beyond prediction, position-resolved flexibility footprints revealed deformation signatures at cognate motifs and flanking regions across diverse TF families. For SOX11, model-derived footprints aligned with DNA shape fluctuations from nanosecond-to-microsecond molecular dynamics trajectories of SOX11-bound nucleosomes, consistent with independently observed DNA conformational dynamics and bound-state stabilization. The in vivo data showed a similar but more context-dependent pattern. OCT4 occupancy tended to correlate with local flexibility, whereas GATA3-pioneered regions showed flexibility coupled with altered rotational positioning of cognate motifs. Flexibility-augmented classifiers further improved discrimination of occupied nucleosomal motifs across ENCODE datasets. Torsional flexibility features, particularly twist dispersion and trx, were most informative for classification.

**Conclusions:** Sequence-derived DNA conformational flexibility provides a quantitative and interpretable representation of sequence context in TF recognition on nucleosomes. By augmenting sequence with structural information, these models help quantify and interpret an indirect-readout contribution in which DNA deformation tendencies may complement motif sequence and DNA shape. This framework may help explain why only selected motif instances are engaged in chromatin, without treating flexibility as independent of primary sequence.

## Background

Transcription factors (TFs) direct gene-regulatory programs by recognizing short DNA sequence motifs embedded in genomic DNA []. In eukaryotes, however, these motifs are interpreted within chromatin, where approximately 75–90% of genomic DNA is packaged into nucleosomes [1]. Each nucleosome wraps ∼147 bp of DNA around a histone octamer in ∼1.7 left-handed superhelical turns, generating a series of histone-contacting superhelical locations (SHLs) arranged around the dyad axis, defined as SHL 0 [2–4]. This organization restricts direct access to many potential cognate motifs and imposes additional constraints arising from steric occlusion, histone-DNA contacts, rotational register, translational position, and local DNA deformation [5,6]. Nucleosomal packaging therefore does not merely obstruct TF binding; it creates a regulatory environment in which the same motif can have different binding outcome depending on its chromatin and sequence context [7].

Pioneer transcription factors (PTFs) are distinguished by their ability to engage target sites in compacted chromatin and promote regulatory states permissive for subsequent factor binding [6,8]. They can act by binding nucleosomal DNA, recruiting chromatin remodelers, destabilizing local histone-DNA contacts, or combining these mechanisms [9–11]. However, the distinction between pioneer and non-pioneer TFs is increasingly viewed as a continuum of “chromatin sensitivity” rather than a strict binary classification [12,13]. Large-scale biochemical assays have shown that many TFs can bind nucleosomal DNA to some extent, whereas in vivo occupancy remains highly selective [14,15]. Because TF motifs are short, degenerate, and widely repeated across eukaryotic genomes [5,7,16], motif identity alone cannot explain why only a subset of potential sites becomes productively occupied in a given cellular context.

Structural studies have begun to reveal how TFs engage motifs within nucleosomal DNA [17,18]. Transcription factors with HMG-domain like SOX2 and SOX11 deform DNA at nucleosomal SHL ±2, widening the minor groove and locally displacing DNA away from the histone surface [19]. OCT4 can bind nucleosomal DNA through its POU-specific domain while other motif elements remain sterically constrained [16,20]. GATA3 can recognize a partial 5^′^-GAT-3^′^ motif presented across solvent exposed major grooves rather than a continuous canonical motif [21]. FOXA1 and GATA4 can reposition enhancer nucleosomes by bending linker DNA and altering histone-DNA contacts [22]. Beyond TFs, nucleosome-associated DNA-binding proteins like UV-DDB can shift nucleosomal DNA to place otherwise occluded substrates into accessible registers [4,23]. Although these mechanisms differ, they allude to a common principle where productive recognition of nucleosomal DNA often requires the local DNA sequence to accommodate bending, twisting, groove remodeling [24]. This makes the mechanical compatibility of a motif and its surrounding sequence a plausible contributor to nucleosomal TF recognition [25].

DNA sequence encodes structural information at several related levels [26,27]. DNA shape describes the preferred geometry of the double helix [28], including minor groove width, propeller twist, roll, helix twist, and electrostatic potential. DNA conformational flexibility, in contrast, refers to the sequence-dependent capacity of DNA to deviate from its preferred geometry through bending, torsional, rotational, or stretching related deformation [29–32]. Shape and flexibility are therefore complementary descriptions of sequence-encoded DNA structure [33]. In free DNA, both contribute to TF specificity through indirect readout, allowing proteins to recognize not only base identity but also the structural and energetic properties of the binding site and its flanks [34–38]. In nucleosomes, these properties become especially relevant because DNA is already sharply bent, rotationally phased, and constrained by histone contacts [3,39]. A TF motif embedded in nucleosomal DNA is therefore evaluated not only as a sequence pattern, but also as part of a mechanically constrained substrate.

The relationship between DNA flexibility and nucleosomal DNA is highly context dependent, and can not be reduced to a single mechanism like ease of DNA wrapping and unwrapping from nucleosome [40]. Sequence encoded flexibility may influence TF engagement through several non exclusive routes. At the nucleosome scale, the broader DNA sequence can affect wrapping, positioning [41], end breathing, local accessibility, and the stability of histone-DNA contacts [42–44]. At a more finer scale, local bending, torsional, rotational, or stretching around the TF binding motifs may influence whether the DNA can adopt a TF-compatible geometry once the motif becomes transiently accessible [45]. Context dependent DNA deformation may facilitate exposure or induced fit [46–48]; in others, mechanically compatible DNA may stabilize the wrapped or TF-bound state [49,50]. Thus, flexibility is best considered as one interpretable representation of sequence context that may contribute to motif exposure, initial engagement, local distortion, or post-binding stabilization [51].

Recent studies support the importance of nucleosomal sequence context in TF binding. NCAP-SELEX provided a large-scale view of TF-nucleosome binding preferences and showed that nucleosomal binding is more widespread than expected from canonical pioneer factors alone [14]. PIONEAR-seq extended this problem by measuring full-length TF binding to reconstituted nucleosome core particles assembled on defined templates, including Widom 601 and genomic nucleosome-positioning sequences [15]. This work showed that local bendability and nucleosomal template sequence influence TF binding positions: TFs classified as dyad or periodic binders on W601 can shift toward end binding on genomic templates, indicating that nucleosomal binding mode is shaped by sequence context rather than being an invariant property of the TF. In vivo, single-molecule and genomic studies further suggest that nucleosomal DNA can contain localized accessible regions associated with TF motifs and TF occupancy [52]. Together, these observations point to sequence-encoded DNA mechanics as a candidate layer of chromatin-sensitive TF recognition.

Despite these advances, several questions remain unresolved. First, it is unclear whether sequence-derived flexibility descriptors quantitatively improve prediction of TF-nucleosome binding affinity across a broad panel of TFs. Next, existing high-throughput nucleosomal studies have mainly emphasized bendability as a measure of cyclizability, whereas diverse form of deformation modes may also known to contribute in nucleosomal TF recognition. Third, the relationship between flexibility descriptors and higher-order sequence composition remains difficult to separate, as such descriptors are themselves computed from nucleotide sequence. Fourth, it remains unclear whether sequence-encoded mechanical signatures can help distinguish occupied nucleosomal binding sites from the larger pool of motif instances that remain unbound in vivo.

Our previous work and related studies have shown that sequence-dependent DNA structural and flexibility descriptors are characteristic and conserved features of cis-regulatory regions across the domains of life [30,53–57]. When incorporated into predictive models, these descriptors have improved the classification of promoter regions from the genomic sequences [58–60]. In particular, conformational flexibility of flanking DNA has been shown to modulate TF binding affinity even when the core motif is unchanged, indicating that the sequence environment surrounding a motif can influence the energetic compatibility of TF-DNA complex formation [37,38]. More recently, flexibility augmented models have improved quantitative prediction of TF binding to free DNA, supporting the idea that sequence-derived flexibility features represent a component of indirect readout that complements nucleotide sequence and DNA shape information [61].

Here, we extend this framework to TF recognition in nucleosomal DNA by developing quantitative models that incorporate sequence-derived DNA flexibility features, building on previous modeling approaches for TF-DNA specificity [36,62]. We curated five DNA flexibility descriptors representing bendability, torsional deformation, backbone conformational variability, and stretching properties, and implemented them in DNAflexpy, a Python package that can be used to transform DNA sequences into position resolved flexibility profiles. We first estimated TF binding affinities to nucleosomal DNA from in vitro NCAP-SELEX experiments [14], and asked whether DNAflexpy-derived descriptors improve TF-nucleosome affinity prediction across 226 datasets representing nearly 150 TFs. Because these descriptors are derived from nucleotide sequence, we do not treat them as independent of primary sequence. Instead, we evaluate whether they provide a compact and physically interpretable representation of sequence context by comparing flexibility-augmented models with mononucleotide, dinucleotide, and trinucleotide sequence baselines, as well as with independently developed representations of DNA conformational fluctuation [63].

We further examine whether flexibility-informed models generalize across different nucleosomal templates, including genomic nucleosome-positioning sequences [15]. To connect predictive performance with structural interpretation, we derive position-resolved flexibility footprints and compare them with the SOX11-DNA experimental structure and all-atom molecular dynamics simulations of SOX-bound nucleosomes [64]. Finally, we integrate ChIP-seq, MNase-seq, and chromatin-accessibility datasets for OCT4, SOX2, KLF4, GATA3, and a broader set of TFs with reported pioneering activity from ENCODE datasets [65] to test whether flexibility signatures can be used for prediction of TF occupied binding sites from unoccupied nucleosomal motifs.

We do not propose DNA flexibility as a determinant separable from primary sequence. Rather, we ask whether sequence-derived flexibility descriptors provide an interpretable feature space that captures deformation tendencies relevant to TF engagement with nucleosomal DNA. This framework allows us to test whether DNA conformational flexibility refines sequence-based models of TF-nucleosome recognition, whether this signal overlaps with or complements independent representations of DNA flexibility, and whether the structural signals in the motif and their flanking sites are associated with productive TF engagement in chromatin.

## Results

### DNAflexpy flexibility features improve TF-nucleosome affinity prediction beyond sequence baselines

We first asked whether sequence-derived DNA flexibility descriptors improve quantitative prediction of transcription factor (TF) binding affinity on nucleosomal DNA. To this end, we generated TF-nucleosome affinity tables from NCAP-SELEX and trained *L*_2_-regularized ridge-regression models evaluated with nested cross-validation. Five DNAflexpy descriptors were used to represent bending-related, torsional, and stiffness-related sequence features [38,61]. Because these descriptors are derived from nucleotide sequence, we did not treat them as physical measurements independent of DNA sequence. Instead, we asked whether they provide an interpretable representation of sequence context that improves prediction beyond standard sequence encodings.

Using mononucleotide one-hot encoding as the initial sequence baseline, adding DNAflexpy features improved model performance for nearly all datasets. Flexibility augmentation increased cross-validated performance in 99.6 of datasets, with a median Δ*R*^2^ of 10.46 over the 1-mer baseline (Wilcoxon signed-rank test, *p* = 2.92 × 10^−40^; **Fig. 1b**). Among the improved datasets, 86.6 gained more than 5 percentage points, 52.6 gained more than 10 percentage points, and only 15 datasets gained more than 20 percentage points. Thus, flexibility descriptors provide information not captured by mononucleotide identity alone.

**Figure 1.**
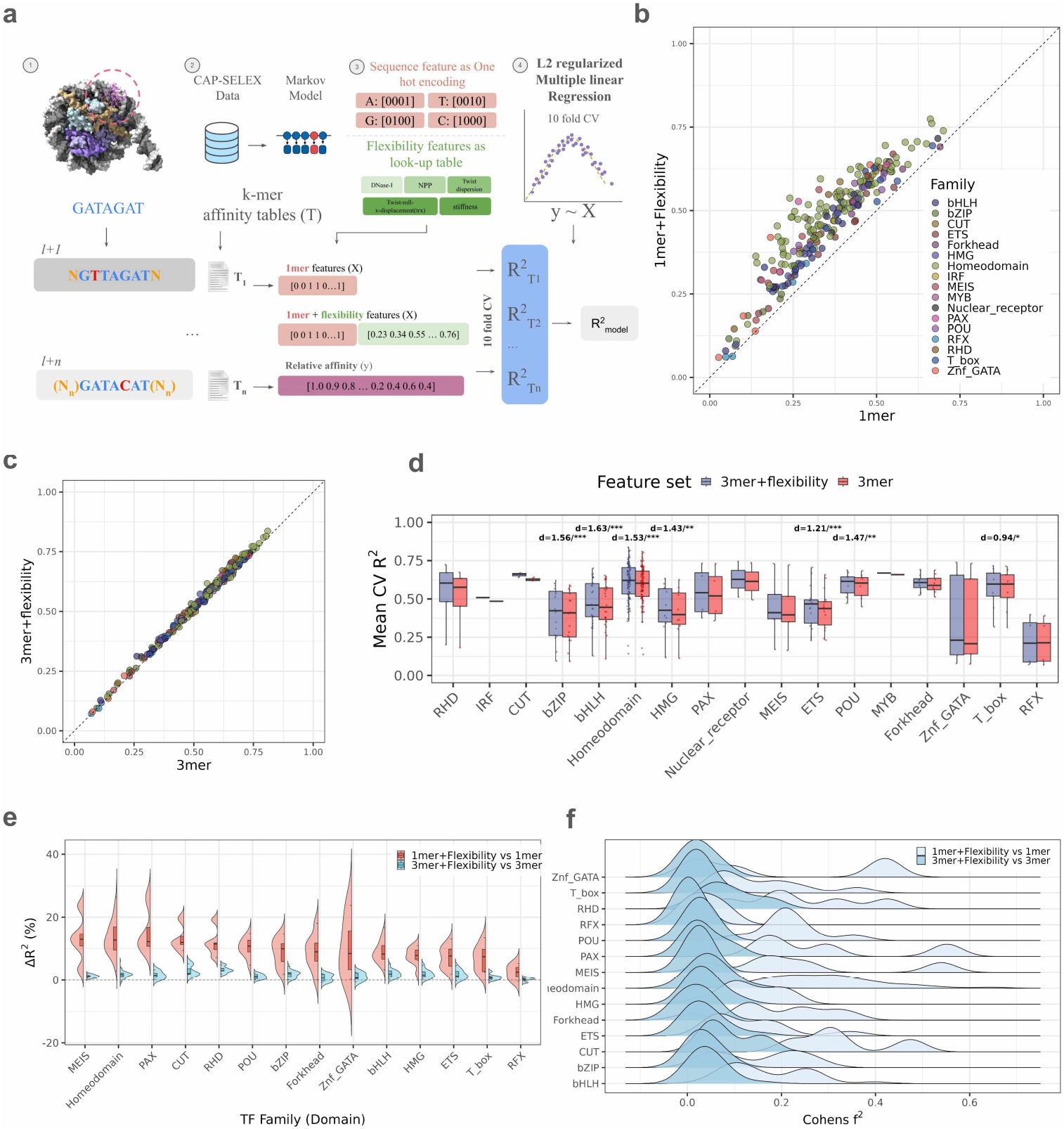
DNA flexibility features improve prediction of TF–nucleosome binding affinity from NCAP-SELEX data. (a) Regression workflow. NCAP-SELEX k-mer count tables were converted to relative TF-nucleosome affinity values; candidate sequences were encoded as sequence features and concatenated with di- and trinucleotide DNAflexpy descriptors to generate the model matrix. *L*_2_-regularized regression models were evaluated under nested cross-validation using 1-mer, flexibility-only, 1-mer + flexibility, 3-mer, and 3-mer + flexibility feature sets. (b) Per-dataset cross-validated *R*^2^ for 1-mer versus 1-mer + flexibility models, colored by DNA-binding domain (DBD) family. Points above the diagonal indicate improved performance after flexibility augmentation (99.6 of datasets; median Δ*R*^2^ = 10.46; Wilcoxon signed-rank test, *p* = 2.92 × 10^−40^). (c) Equivalent comparison for 3-mer versus 3-mer + flexibility models. Flexibility retains residual predictive value after trinucleotide composition is modelled (212/226 datasets above the diagonal; median Δ*R*^2^ = 1.67; Wilcoxon signed-rank test, *p* = 1.52 × 10^−36^). (d) Family-stratified mean cross-validated *R*^2^ for 3-mer and 3-mer + flexibility models. Family-level Δ*R*^2^ differences remain significant (Kruskal-Wallis test, *H* = 25.25, *p* = 4.89 ×10^−3^), with per-family Wilcoxon significance indicated above paired comparisons. (e) Split-density distributions of Δ*R*^2^ for flexibility augmentation over the 1-mer baseline and over the 3-mer baseline. PAX, Homeodomain, CUT, MEIS, and RHD show the largest gains over 1-mer models, whereas RHD, CUT, bHLH, and HMG retain the largest residual gains over 3-mer models. (f) Cohen’s *f* ^2^ effect-size densities by family. Most families exceed the medium-effect threshold over the 1-mer baseline, led by CUT (*f* ^2^ = 0.33), PAX (0.30), Homeodomain (0.29), and RHD (0.24). Over the 3-mer baseline, RHD, CUT, Homeodomain, and bHLH exceed the small-effect threshold (*f* ^2^ > 0.02).

We next tested whether these gains could be explained by higher-order sequence composition. Dinucleotide and trinucleotide encodings capture nearest-neighbour dependencies, local stacking context, and part of the same sequence information from which several flexibility descriptors are derived. As expected, 2-mer and 3-mer models reduced the apparent gain from flexibility. The 1-mer + flexibility models performed comparably to the larger 2-mer models, whereas 3-mer models clearly outperformed both (**Fig. S1**). Flexibility-only models performed below the *k*-mer baselines, consistent with the view that flexibility descriptors refine, but do not replace, local sequence information.

We also evaluated whether flexibility retained predictive value after trinucleotide sequence context had already been included. Adding flexibility descriptors to the 3-mer baseline produced positive Δ*R*^2^ in 212 of 226 datasets (93.8; Wilcoxon signed-rank test, *p* = 1.52 × 10^−36^), with a median improvement of 1.67 (**Fig. 1c**). This residual gain was smaller than the gain observed over the 1-mer baseline, but it indicates that the flexibility descriptors capture predictive information not fully represented by trinucleotide composition.

The contribution of flexibility varied across DNA-binding domain (DBD) families. Over the 1-mer baseline, the largest mean gains were observed for PAX, Homeodomain, CUT, MEIS, and RHD families, with mean Δ*R*^2^ values of 15.29, 13.41, 13.22, 12.67, and 11.88, respectively. RFX showed the smallest mean gain (2.48). These family-level differences were statistically significant (Kruskal-Wallis test, *H* = 50.53, *p* = 2.13 × 10^−7^; **Fig. 1d**). Over the 3-mer baseline, absolute gains were smaller but family-level heterogeneity remained detectable (Kruskal-Wallis test, *H* = 25.25, *p* = 4.89 × 10^−3^; **Fig. 1e**). RHD, CUT, bHLH, and HMG retained the largest mean positive gains over 3-mer models, with mean Δ*R*^2^ values of 3.28, 2.91, 2.02, and 1.91, respectively. Forkhead factors, in contrast, gained 8.84 over the 1-mer baseline but showed a small negative mean gain over the 3-mer baseline (−0.53), suggesting that much of the flexibility-associated signal for this family is already represented by trinucleotide composition.

Cohen’s *f* ^2^ effect sizes, computed to quantify the residual improvement of the flexibility-augmented model over the sequence baseline, yielded the same conclusion (Methods; **Fig. 1f**). Over the 1-mer baseline, most TF families exceeded the medium-effect threshold (0.15), with the largest effects observed for CUT (*f* ^2^ = 0.33), PAX (*f* ^2^ = 0.30), Homeodomain (*f* ^2^ = 0.29), and RHD (*f* ^2^ = 0.24). Over the 3-mer baseline, effect sizes decreased, but RHD (*f* ^2^ = 0.091), CUT (*f* ^2^ = 0.085), Homeodomain (*f* ^2^ = 0.046), and bHLH (*f* ^2^ = 0.041) still exceeded the small-effect threshold (0.02). Forkhead and RFX showed negligible or slightly negative effects. Together, these analyses show that DNAflexpy features provide the strongest improvement when the sequence baseline is simple, but also retain a smaller residual contribution after local trinucleotide context is modelled. This residual signal is family-dependent rather than universal.

### DNAflexpy and Deep DNAshape identify overlapping structural signals in TF-nucleosome binding

We next asked whether the predictive signal captured by DNAflexpy reflects a broader sequence-encoded structural signal or is specific to the parametrization of DNAflexpy descriptors. To test this, we compared DNAflexpy with shape-fluctuation features predicted by Deep DNAshape [63]. Deep DNAshape estimates sequence-dependent fluctuations in DNA shape parameters using a learned representation that incorporates extended flanking sequence context, whereas DNAflexpy encodes empirical descriptors derived from DNase I cleavage preferences, nucleosome-positioning preferences, and structural measurements of DNA deformation. The two approaches therefore differ in origin, feature formulation, and modelling of sequence context.

Despite these differences, DNAflexpy and ShapeFL showed strong concordance across the NCAP-SELEX panel (**Fig. 2a,b**). DNAflexpy marginally outperformed ShapeFL features when added to 1-mer models (**Fig. 2c**). Per-dataset Δ*R*^2^ values for DNAflexpy and ShapeFL-L0 were strongly correlated over both the 1-mer and 3-mer baselines (Spearman *ρ* = 0.933 and *ρ* = 0.742, respectively; **Fig. S2a**), indicating that the two encodings identified largely overlapping datasets in which structural information improved prediction. Against the stronger 3-mer baseline, the deeper ShapeFL-L2 layer retained a slight advantage in some comparisons (**Fig. 2d**), suggesting that learned fluctuation features from additional sequence context may capture relevant signal when trinucleotide composition is already represented.

**Figure 2.**
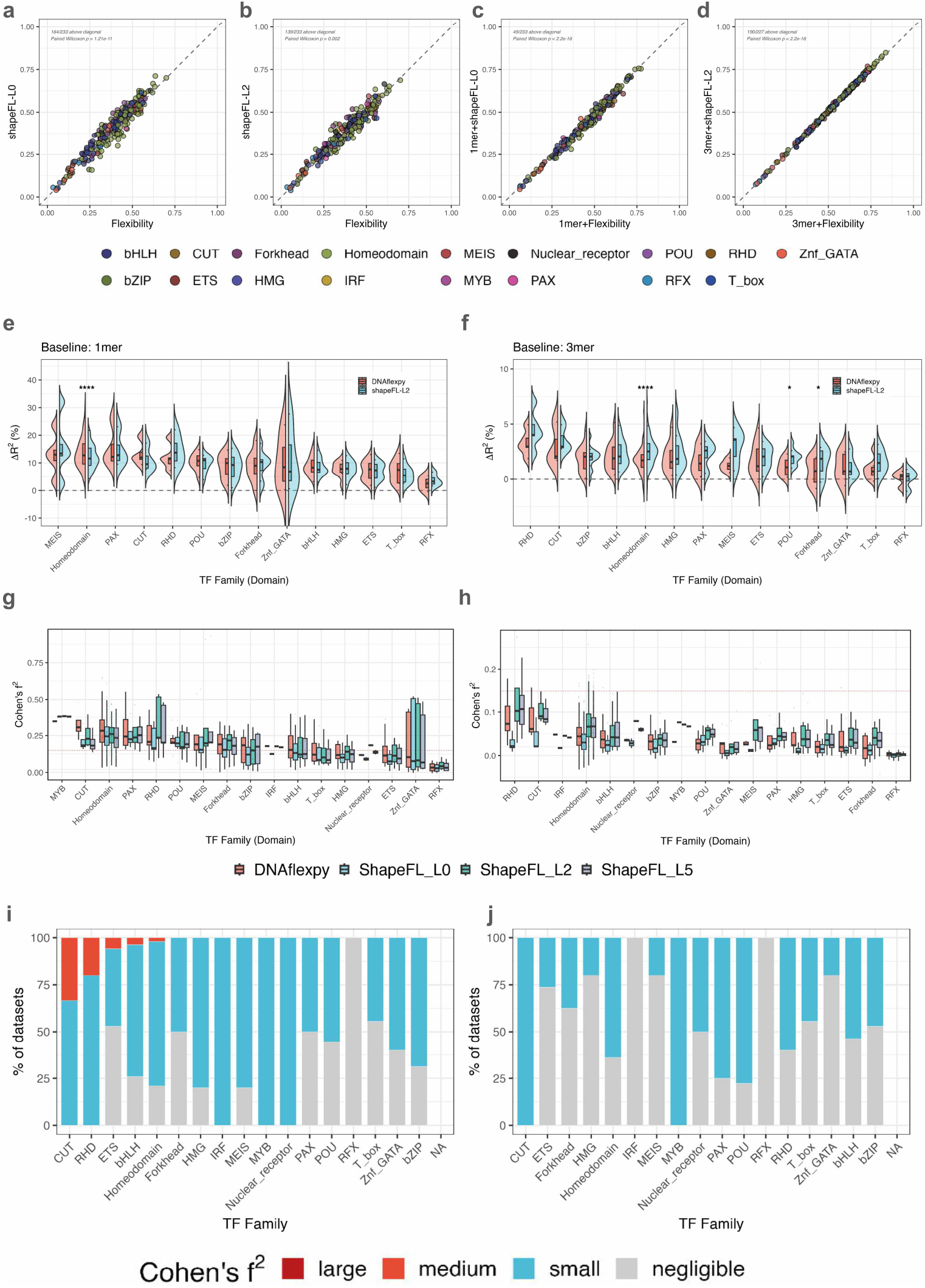
DNAflexpy descriptors recover binding-relevant structural signal comparable to DeepDNAshape shape-fluctuation features. (a) Per-dataset performance for flexibility-only versus ShapeFL-L0-only models across NCAP-SELEX datasets, colored by DBD family. (b) Equivalent comparison using ShapeFL-L2, which incorporates extended flanking sequence context. (c) Per-dataset *R*^2^ for 1-mer + DNAflexpy versus 1-mer + ShapeFL-L0 models. The two structural encodings track each other closely, with a small overall advantage for DNAflexpy in this comparison. (d) Per-dataset *R*^2^ for 3-mer + DNAflexpy versus 3-mer + ShapeFL models. Against the stronger trinucleotide baseline, ShapeFL shows a small advantage in some datasets. Per-dataset Δ*R*^2^ values for DNAflexpy and ShapeFL-L0 are strongly concordant over both the 1-mer and 3-mer baselines (Spearman *ρ* = 0.933 and *ρ* = 0.742, respectively; Fig. S2a). (e) Family-stratified Δ*R*^2^ over the 1-mer baseline for DNAflexpy and ShapeFL-L2. MEIS, Homeodomain, PAX, CUT, and RHD show larger gains, whereas ETS, T-box, and RFX show weaker gains. (f) Equivalent comparison over the 3-mer baseline. Gains are compressed below approximately 5 for most families, and family-level ordering is partly preserved. (g) Cohen’s *f* ^2^ over the 1-mer baseline for DNAflexpy, ShapeFL-L0, ShapeFL-L2, and ShapeFL-L5. (h) Equivalent comparison over the 3-mer baseline; DNAflexpy remains slightly ahead of ShapeFL-L0, whereas deeper ShapeFL layers add limited additional signal in this regression framework. (i) Per-family fraction of TF datasets in each Cohen’s *f* ^2^ category for DNAflexpy added to the 3-mer baseline. (j) Equivalent breakdown for ShapeFL-L0 added to the 3-mer baseline. The higher-effect categories are more frequently populated by DNAflexpy.

Family-stratified performance comparisons reinforced this convergence. Over the 1-mer baseline, both DNAflexpy and ShapeFL-L2 produced positive Δ*R*^2^ values across most TF families, with stronger gains for PAX, CUT, POU, and Homeodomain factors and weaker gains for ETS, T-box, and RFX factors (**Fig. 2e**). Over the 3-mer baseline, gains dropped below approximately 5 for most families, and the family-level ordering became flatter (**Fig. 2f**). Thus, the family ordering in the comparison was modestly preserved across feature types, but compressed by the richer sequence baseline.

Effect-size analyses showed the same pattern. Across most families, DNAflexpy performed comparably to deeper ShapeFL layers that incorporate longer sequence context (**Fig. 2g,h**). However, increasing ShapeFL context depth to L2 or L5 produced no additional benefit in this regression framework across the current datasets (**Fig. S2b**). Within TF families, datasets with higher *f* ^2^ values over the 3-mer baseline were more frequently enriched for DNAflexpy than for ShapeFL-L0 features (**Fig. 2i,j**). These results suggest that DNAflexpy and ShapeFL recover mostly overlapping but non-redundant structural information and may represent partly distinct aspects of sequence-dependent conformational information. The convergence with an independent shape-fluctuation model argues against the flexibility signal being an artifact of a single parametrization.

### Flexibility associated predictive gain in specific to TFs and nucleosomal templates

To test whether flexibility-associated gains generalize beyond a single assay, we also analysed PIONEAR-seq datasets [15]. Although NCAP-SELEX provides a large-scale estimate of TF binding affinity on nucleosomal DNA [14], the assay is initialized from randomized sequence pools and is therefore less suited for testing how TF engagement varies across defined nucleosomal templates. Mariani *et al*. tiled 20-bp randomized segments across W601 and three genomic positioning sequences, ALB1, CX3, and NRCAM, allowing template-dependent TF-nucleosome affinity to be modelled directly.

Applying the same ridge-regression framework to PIONEAR-seq datasets revealed a consistent hierarchy of model performance across the assayed TF and template combinations (**Fig. 3a**). Flexibility-only models matched or exceeded 1-mer models for several combinations, indicating that sequence-derived flexibility descriptors alone capture part of the affinity landscape. Adding flexibility to the 1-mer encoding improved mean cross-validated *R*^2^ for most TFs. As expected, 3-mer models outperformed 1-mer models, and the 3-mer + flexibility model gave the strongest overall performance. Mean *R*^2^ increased from 0.302 for 1-mer and 0.291 for flexibility-only models to 0.374 for 1-mer + flexibility. While 3-mer models performed best among the sequence-only models (*R*^2^ = 0.497), 3-mer + flexibility further improved performance (*R*^2^ = 0.507). Thus, flexibility features augmented sequence-based prediction, indicating that sequence-only models do not fully capture TF binding affinity in nucleosomal context.

**Figure 3.**
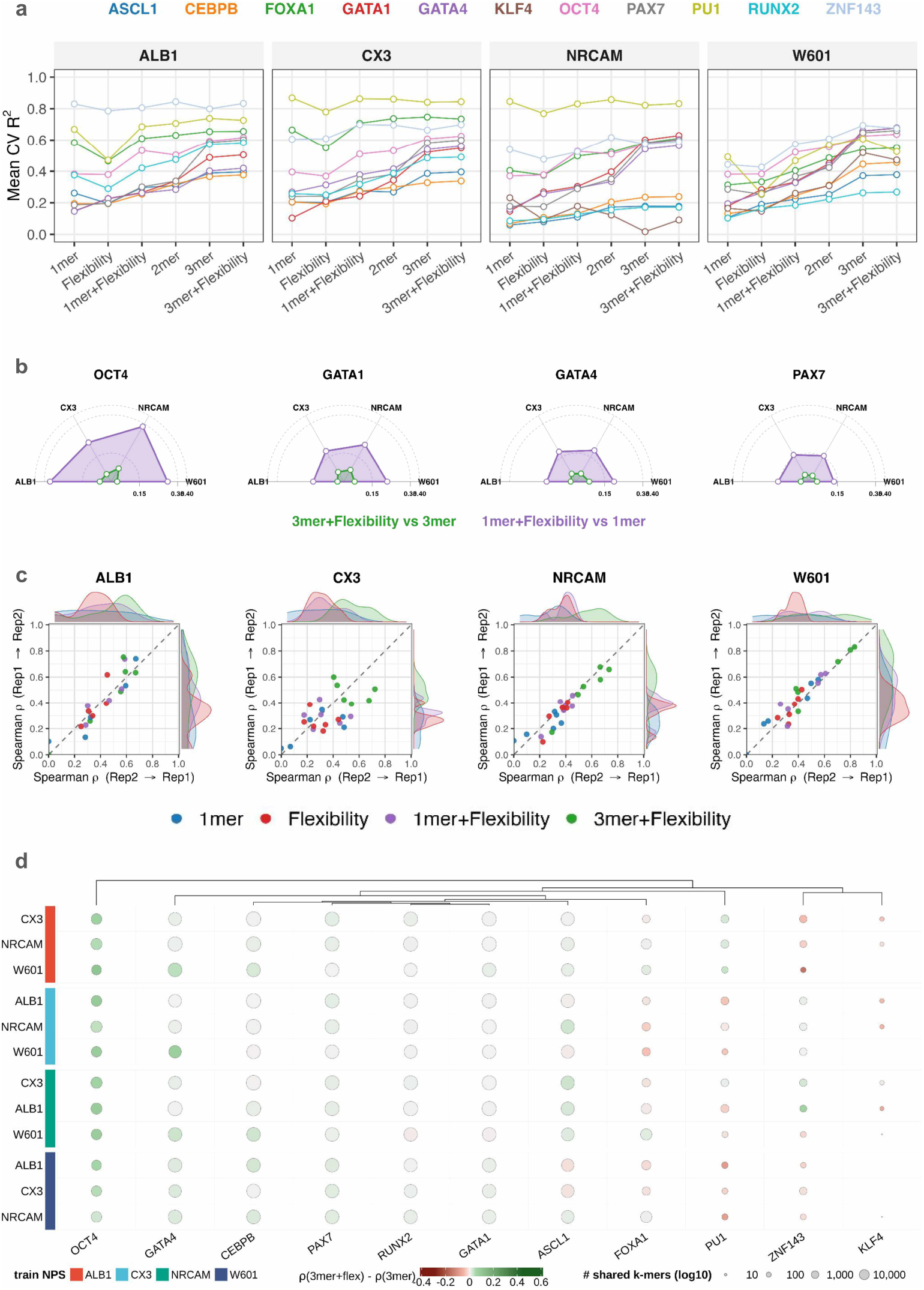
DNAflexpy generalizes to PIONEAR-seq and tracks template-dependent TF–nucleosome binding. (a) Mean 10-fold cross-validated *R*^2^ for *L*_2_-regularized models across the four PIONEAR-seq templates: ALB1, CX3, NRCAM, and W601. Each TF is shown under 1-mer, flexibility-only, 1-mer + flexibility, 3-mer, and 3-mer + flexibility encodings. Mean *R*^2^ across TF-template combinations increases from 0.302 for 1-mer and 0.291 for flexibility-only models to 0.374 for 1-mer + flexibility, 0.497 for 3-mer, and 0.507 for 3-mer + flexibility. (b) Per-TF radar plots for representative TFs. Each vertex represents a nucleosomal template. Polygons compare flexibility-augmented models against their sequence-only baselines, showing template-dependent gains. (c) Cross-replicate concordance across PIONEAR-seq templates. Each panel compares predictions from reciprocal train-test replicate evaluations, with points colored by model class. Replicate stability increases from *ρ* = 0.260 for 1-mer models to *ρ* = 0.520 for 3-mer + flexibility models. (d) Cross-template generalization matrix. Rows represent directed train-to-test template pairs grouped by training template, and columns represent TFs. Bubble color reports the performance change from adding flexibility to the 3-mer baseline, and bubble size reports the number of shared k-mers between training and test sets on a log_10_ scale. OCT4, GATA4, CEBPB, and PAX7 show predominantly positive cross-template gains, whereas PU.1, ZNF143, and KLF4 show weaker or negative effects.

The magnitude of flexibility-associated gain differed across nucleosomal templates. Gains over the 3-mer baseline were highest in NRCAM and lowest in W601, indicating that the predictive value of flexibility depends on the underlying nucleosome-forming sequence. OCT4, GATA1, GATA4, and PAX7 showed clear flexibility-associated gains across multiple templates, with particularly strong effects over the 1-mer baseline for OCT4 and the GATA factors (**Fig. 3b**; **Fig. S3a**). These gains were not distributed uniformly across ALB1, CX3, NRCAM, and W601, consistent with the central conclusion of PIONEAR-seq that TF binding propensities depend on nucleosomal template sequence. Flexibility-associated improvements were larger over the simple 1-mer baseline and more context-specific over the 3-mer baseline.

The predictive signal from flexibility features was also reproducible across biological replicates. When models were trained on one biological replicate and evaluated on the other, the Spearman correlation between predicted and observed binding increased with feature richness (**Fig. 3c**). Mean cross-replicate concordance improved from *ρ* = 0.260 for 1-mer models and *ρ* = 0.314 for flexibility-only models to *ρ* = 0.373 for 1-mer + flexibility and *ρ* = 0.520 for 3-mer + flexibility. Template identity also shaped feature performance: FOXA1, GATA1, GATA4, OCT4, PAX7, and RUNX2 differed in whether their strongest performance was observed on W601 or on genomic templates. Across templates, 3-mer + flexibility produced the most reproducible profiles, whereas 1-mer and flexibility-only models were more dispersed (**Fig. S3c**).

Because predictive performance was influenced by nucleosomal templates, we finally tested whether flexibility-augmented models trained on one nucleosomal template could generalize to other templates. For each TF, models were trained on one template and evaluated on affinity data from the remaining three templates. Cross-template evaluation revealed predominantly positive flexibility effects above the 3-mer baseline for OCT4, GATA4, CEBPB, PAX7, and several other TFs, whereas PU.1, ZNF143, and KLF4 showed weaker or negative effects (**Fig. 3d**). Thus, DNAflexpy features represent TF-specific conformational preferences that are not purely template-specific; rather, transferability depends on both TF identity and nucleosomal sequence background.

Together, NCAP-SELEX and PIONEAR-seq show that flexibility-associated gains are reproducible across assay designs but modulated by TF identity, template sequence, and baseline complexity. This supports a model in which nucleosomal TF affinity is shaped by deformation-related properties of the surrounding sequence in addition to motif identity.

### Position resolved flexibility footprints recover TF mediated DNA deformation signatures

The improved predictive performance of DNAflexpy descriptors across large-scale TF-nucleosome affinity datasets prompted us to ask whether flexibility-augmented models could also reveal where flexibility contributes to structural readout [66–68]. We therefore systematically interpreted the cross-validated models and focused on SOX/HMG factors, whose DNA-binding domains bend and deform DNA, providing a well-defined structural benchmark [69,70]. In the SOX-bound nucleosome structure reported by Dodonova *et al*., the HMG domain recognizes a 5^′^-TTGT-3^′^ motif near SHL ±2 and induces local DNA deformation [19]. We extracted position-specific “flexibility footprints” from flexibility-augmented models trained on in vitro affinity datasets and compared them with three independent structural references: the SOX11-DNA crystal structure, affinity-stratified DNA shape profiles, and all-atom molecular dynamics (MD) simulations of SOX-bound and SOX-unbound nucleosomes.

Flexibility footprints were derived using a position-specific model-interpretation strategy adapted from DNA shape-readout analyses [34,36,61]. For each TF, we trained a 1-mer baseline model with performance 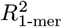, added flexibility descriptors for one nucleotide position *i* at a time, and estimated 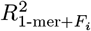. The position-specific gain was 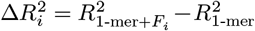, and the relative contribution was 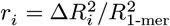. Plotting *r*_*i*_ across the binding site defines a TF-specific flexibility footprint that estimates where flexibility contributes most strongly to affinity prediction.

For SOX TFs, these footprints peaked within the motif and extended into adjacent flanking positions (**Fig. 4a**). In the SOX11-DNA crystal structure, Phe56 and Met57 insert into the minor groove, while Asn54 supports localized bending near the central 5^′^-TG-3^′^ step of the 5^′^-TTGT-3^′^ motif (**Fig. 4b**). The footprint corresponding to the 5^′^-ACAA-3^′^ representation in **Fig. 4a** therefore maps to the same motif region that undergoes direct structural deformation in the SOX11-DNA complex.

**Figure 4.**
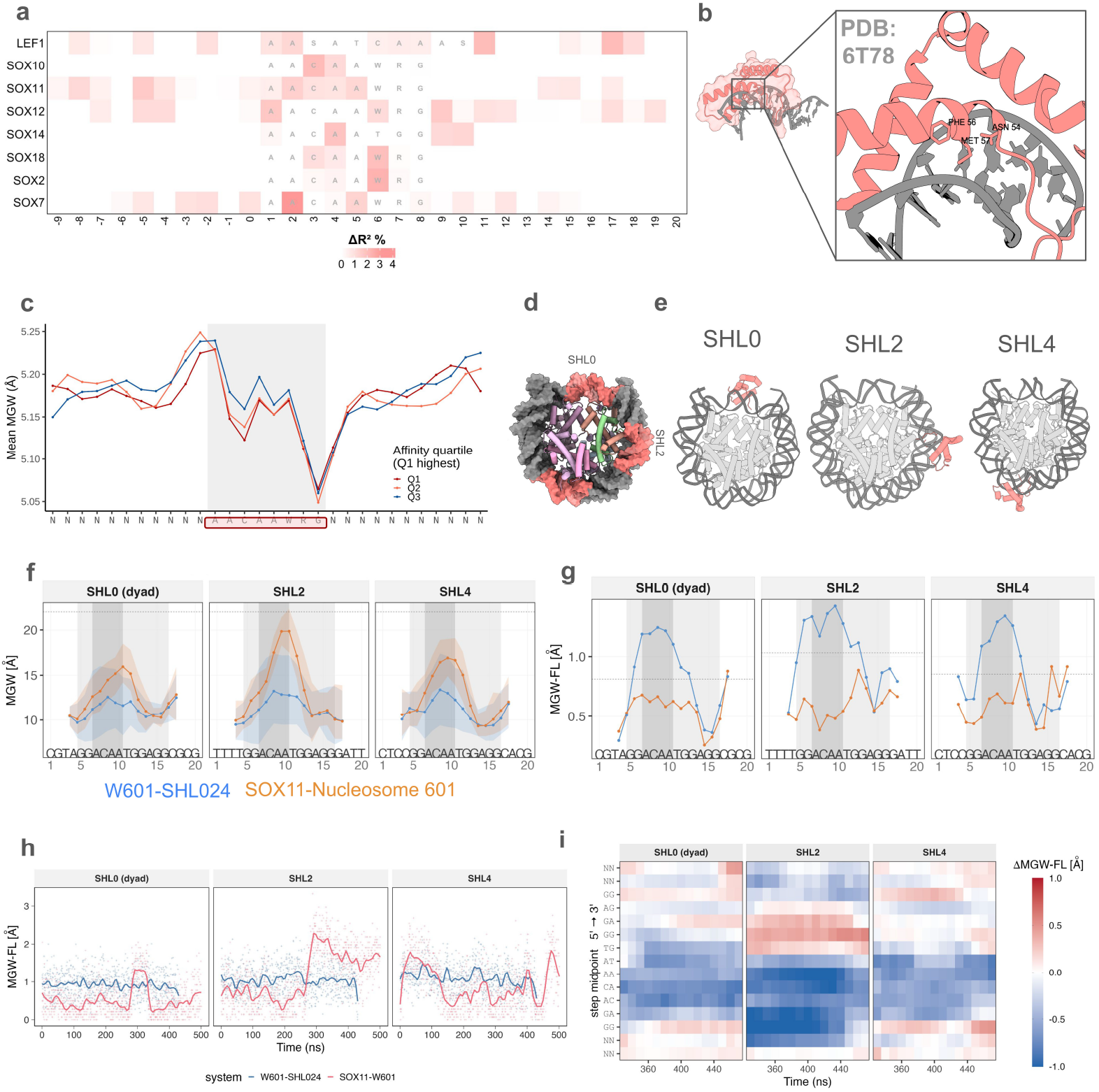
Model-derived flexibility footprints match structural and atomistic measures of SOX–nucleosome indirect readout. (a) Position resolved flexibility footprint(contribution) heatmap for TFs with HMG-domain. Rows represent TFs, and columns represent nucleotide positions across the aligned binding site. Color encodes the per-position flexibility contribution, 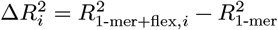, normalized to the 1-mer baseline. Motifs recovered for each TF from NCAP-SELEX study were overlayed in the heatmap. (b) SOX11-DNA structure (PDB ID: 6T78). Phe56 and Met57 insert into the minor groove, whereas Asn54 contacts the central 5^′^-TG-3^′^ step of the 5^′^-TTGT-3^′^ motif. (c) Average minor groove width (MGW) profiles at SOX binding sites, stratified by NCAP-SELEX affinity. Q1 represents the highest-affinity sites and Q3 the lowest-affinity sites. The largest MGW differences localize to the central TG/CA step and adjacent flanks. (d) SOX motif insertion design on widom 601 template for MD simulations on mutated W601 nucleosomal templates by Ozden et al. (e) Nucleosome cartoons showing the inserted SOX motif at SHL0, SHL2, or SHL4 by same study. (f) MGW across the inserted SOX motif for SOX-unbound (W601-SHL024), and SOX11-bound trajectories at SHL0, SHL2, and SHL4. Minor-groove widening upon SOX binding is strongest at SHL2. (g) Per-position MGW fluctuation (MGW-FL) for the same simulations, used as a local measure of conformational fluctuation. The SHL2 SOX core shows higher pre-binding MGW-FL than SHL0 or SHL4. (h) Time-resolved MGW at the central TG/CA step for unbound and bound nucleosomes at SHL0, SHL2, and SHL4. The bound SHL2 trajectory samples a sustained wide-groove state, whereas SHL0 and SHL4 do not show the same persistent bound-like geometry. (i) Δ MGW-FL heatmap, calculated as bound minus unbound fluctuation across motif positions and time. SHL2 shows the strongest localized post-binding reduction in MGW fluctuation around the motif core.

Affinity-stratified DNA shape profiles supported this mapping. NCAP-SELEX sites were grouped into Q1 high-affinity and Q3 low-affinity classes. MGW profiles showed that high-affinity sequences had narrower groove width than low-affinity sequences, with the clearest difference around the central 5^′^-TG/CA-3^′^ step (**Fig. 4c**). Thus, the positions where flexibility most improved prediction were aligned with the positions where DNA shape separated high-from low-affinity SOX sites.

We then asked whether conformational dynamics at these positions explain preferential SOX binding in nucleosomes. Replicated MD trajectories were analysed for SOX-unbound nucleosomes carrying SOX motifs at SHL0, SHL2, and SHL4 (**Fig. 4d**) and for SOX-bound nucleosomes at the same locations (**Fig. 4e**). Previous work showed that SOX binding is favoured at SHL2, weaker at SHL4, and disfavoured at the dyad [64]. This positional preference was attributed to histone-DNA clamping at the dyad and to a steric clash between H2A and SOX at SHL4. SHL2 lacks both constraints, allowing the minor-groove widening required for stable SOX engagement.

SOX-unbound nucleosomal motifs rarely sampled the bound-like minor-groove geometry, even at productive SHL2 (**Fig. 4f**; **Fig. S4a**). Motifs at SHL2 instead showed higher local MGW fluctuation (MGW-FL) around the motif than SHL0 or SHL4 (**Fig. 4g**; **Fig. S4g**,**h**), indicating a broader pre-binding conformational ensemble. This elevated plasticity was localized to the motif core and immediate flanks, matching the footprinted region.

Upon SOX binding, the SHL2 motif adopted and stabilized the widened minor-groove state characteristic of HMG-domain engagement (**Fig. 4h**). Time-resolved MGW profiles showed a pronounced wide-groove state at the central AC/TG step in the bound SHL2 trajectory, whereas SHL0 and SHL4 did not show the same persistent bound-like geometry (**Fig. S4b**). Slide and roll profiles also differed across SHLs, with the largest bound-state perturbations concentrated around the motif core and flanks (**Fig. S4c**,**d**). SOX binding therefore remodels a local DNA-shape ensemble rather than shifting groove width alone.

Finally, comparing bound and unbound trajectories showed that MGW-FL decreased after binding, with the strongest and most localized reduction at SHL2. The time-resolved ΔMGW-FL heatmap revealed a localized reorganization around the SOX motif core, with the strongest negative shift at SHL2 during the later simulation window (**Fig. 4i**). SHL0 and SHL4 showed weaker or more dispersed changes. SHL2 therefore combines elevated pre-binding plasticity with post-binding stabilization of a widened, SOX-compatible minor-groove state.

The SOX analyses converge across model, structure, affinity stratification, and dynamics. The same central motif positions carry the strongest flexibility footprint, undergo structural deformation in the SOX11-DNA complex, separate high-from low-affinity nucleosomal binding sites, show elevated SOX-unbound MGW-FL at SHL2, and stabilize after SOX binding. Because the regression model was not trained on structural or MD information, this agreement supports flexibility footprints as interpretable, position-resolved signatures of indirect readout. It does not prove that flexibility alone causes binding; rather, it links model-derived flexibility contributions to independently observed deformation, pre-binding plasticity, and stabilization upon TF binding.

### Characteristic DNA flexibility signal are detectable at TF occupied in vivo nucleosomal binding sites

We next asked whether the convergent predictive and structural signals observed in prior analyses are detectable at genomic binding sites of TFs known to engage chromatin. We analysed two datasets: OCT4, SOX2, and KLF4 (OSK hereafter) binding in IMR90 cells, and GATA3 binding after ectopic expression in MDA-MB-231 cells.

For OSK factors, whose pioneering activity is established in the literature [71–73], we integrated ChIP-seq and MNase-seq data from IMR90 cells [16] and classified binding sites by local nucleosome occupancy (**Fig. 5a**). DNAflexpy profiles differed between nucleosomal and non-nucleosomal sites, with the strongest contrasts centered on the binding site and immediate flanks, indicating that the in vitro flexibility signal is also detectable in genomic sequence context.

**Figure 5.**
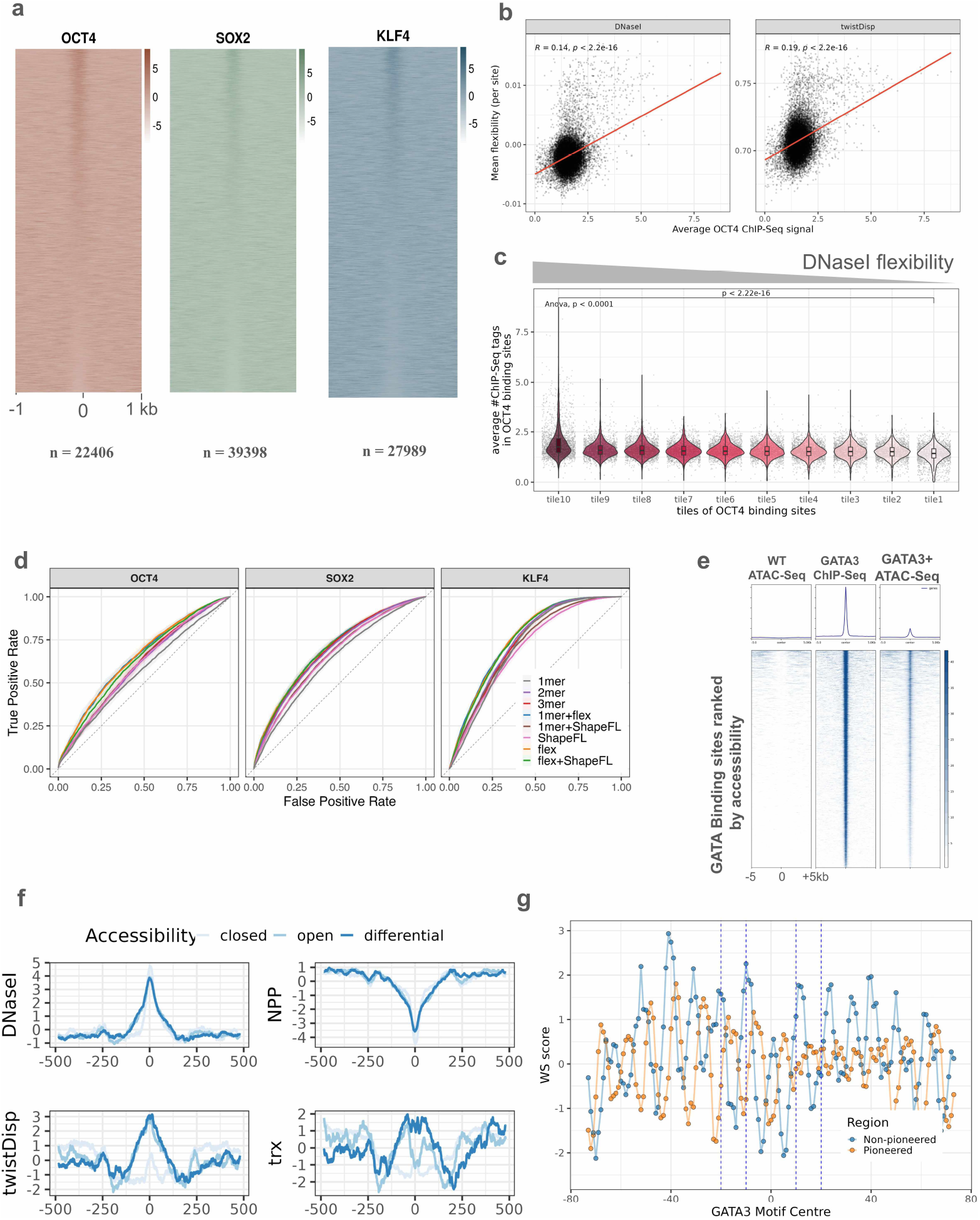
DNA flexibility signatures track pioneer-factor engagement in vivo. (a) OCT4, SOX2, and KLF4 ChIP-seq read-density heatmaps centered on ChIP-seq peak summits in IMR90 cells (±1 kb). Rows are sorted by central MNase-seq signal, and sample sizes are shown below each panel. (b) OCT4 ChIP-seq signal versus local DNA flexibility at nucleosomal OCT4 binding sites for DNase I sensitivity and twist dispersion. Lines show linear fits; partial Spearman correlations after GC-content correction are *ρ* = 0.150 for DNase I sensitivity and *ρ* = 0.222 for twist dispersion. (c) OCT4 ChIP-seq signal at nucleosomal binding sites stratified into deciles of mean DNase I bendability. Decile 10 represents the highest-flexibility group and decile 1 the lowest-flexibility group. Kruskal-Wallis test: *H* = 574.0, *p* = 7.8 × 10^−118^; decile 10 versus decile 1: Cohen’s *d* = 0.73 (95% CI, 0.66 to 0.79). (d) Post hoc ROC evaluation using predicted MNase-seq signal from cross-validated XGBoost regression models to distinguish OSK binding sites in nucleosomal and non-nucleosomal contexts. Feature sets include sequence-only, flexibility-only, ShapeFL-only, and combined structural encodings. (e) GATA3-bound regions in MDA-MB-231 cells ranked by ATAC-seq accessibility. Heatmaps show WT ATAC-seq, GATA3 ChIP-seq, and ATAC-seq after ectopic GATA3 expression. Regions are partitioned into constitutively open, pioneered, and non-pioneered/closed classes. (f) Average DNAflexpy profiles for DNase I sensitivity, NPP, twist dispersion, and trx across GATA3 sites classified by accessibility outcome in a ±500 bp window. Pioneered sites show elevated flanking flexibility relative to closed sites, primarily outside the motif core. (g) Average WW/SS rotational-positioning score from the Cui-Zhurkin model, centered on the GATA3 motif. Pioneered and non-pioneered sites separate, consistent with differential rotational motif presentation on the nucleosome surface.

Because nucleosomal and non-nucleosomal sites may differ in sequence composition, we next tested whether the flexibility profiles could be explained by GC content or dinucleotide composition. OCT4 showed no significant GC-content difference between nucleosomal and non-nucleosomal sites (mean GC, 40.5 versus 40.9; Wilcoxon rank-sum test, *p* = 0.528; Cliff’s *δ* = 0.004). SOX2 showed a small GC-content shift (34.8 versus 33.8; Cliff’s *δ* = 0.109), whereas KLF4 showed a larger difference (45.8 versus 53.1; Cliff’s *δ* = 0.380), consistent with its GC-rich motif preference and binding to promoter-proximal regions. Dinucleotide-preserving sequence shuffling using biasaway (*k* = 2) eliminated the flexibility peaks centered at the binding site (**Fig. S6b**,**c**), suggesting that the observed profiles reflect position-specific sequence organization rather than GC content or dinucleotide composition alone.

Among nucleosomal OCT4 sites, ChIP-seq occupancy increased monotonically across deciles ranked by DNase I bendability or twist dispersion (**Fig. 5c**). Both descriptors showed significant variation across deciles (DNase I sensitivity, *H* = 574.0, *p* = 7.8 × 10^−118^, *η*^2^ = 0.030; twist dispersion, *H* = 865.7, *p* = 1.5 × 10^−180^, *η*^2^ = 0.046; Kruskal-Wallis test). The highest- and lowest-flexibility deciles were separated by moderate-to-large effects, with Cohen’s *d* = 0.73 (95% CI, 0.66 to 0.79) for DNase I bendability and *d* = 0.85 (95% CI, 0.78 to 0.91) for twist dispersion. After controlling for GC content, the partial Spearman correlation increased from *ρ* = 0.141 to *ρ* = 0.150 for DNase I sensitivity and from *ρ* = 0.192 to *ρ* = 0.222 for twist dispersion (**Fig. 5b**). GC content therefore contributes to sequence variation but does not explain the association between flexibility and OCT4 occupancy.

DNA sequence can influence both nucleosome positioning and TF engagement at cis-regulatory elements [74]. We therefore asked whether sequence-derived flexibility features predict the nucleosomal context of OSK binding sites by modelling MNase-seq signal. XGBoost regression models trained on flexibility features outperformed *k*-mer sequence baselines across OCT4, SOX2, and KLF4, reaching *R*^2^ = 0.22/0.16/0.15 and Spearman *ρ* = 0.32/0.41/0.51, respectively. Pooled across TFs, flexibility gained Δ*R*^2^ = +0.079 and Δ*ρ* = +0.134 over the 1-mer baseline (Wilcoxon signed-rank test on 30 outer-fold observations, *p* < 2 × 10^−9^), and remained significant over the 2-mer and 3-mer baselines (Δ*R*^2^ = +0.039 and +0.052; Δ*ρ* = +0.057 and +0.061; all *p* < 6 × 10^−9^). In contrast, ShapeFL did not improve *R*^2^ over the 2-mer or 3-mer baselines (Δ*R*^2^ = −0.030 and −0.017, respectively). As a post hoc analysis, we used the predicted MNase-seq signal from the regression models as a ranking score to distinguish OSK binding sites in nucleosomal and non-nucleosomal contexts (**Fig. 5d**). This evaluation recapitulated the regression performance pattern (Methods), suggesting that DNAflexpy flexibility features provide modest but measurable predictive information for nucleosome-associated OSK binding beyond local *k*-mer composition.

We then examined GATA3, a zinc-finger pioneer factor with known activity across chromatin contexts [21,75–78]. GATA3-bound regions in MDA-MB-231 cells were classified as pioneered, constitutively open, or closed/non-pioneered using ATAC-seq changes after ectopic GATA3 expression (**Fig. 5e**; Methods). GATA motifs occurred across all three classes, but motif presence and motif strength did not fully explain accessibility gain.

Differences in flexibility profiles were observed across these classes. Pioneered regions showed higher DNase I bendability than closed sites, with the difference centered near the motif and its flanks (**Fig. 5f**). Twist dispersion and trx profiles showed related differences, indicating that the sequence context surrounding the motif varies with accessibility outcome. These observations prompted us to ask whether pioneered and non-pioneered regions also differ in nucleosomal rotational positioning, which could alter how the GATA motif is presented on the nucleosome surface. Using the Cui-Zhurkin model [79,80], we found that GATA motifs in pioneered sites showed a distinct WS-score profile compared with non-pioneered GATA-binding sites (**Fig. 5g**), consistent with altered groove presentation relative to the histone surface. Together, these results suggest that GATA3-mediated accessibility gain is associated not only with motif presence, but also with flanking bendability, torsional flexibility, and rotational presentation of the motif-proximal DNA.

Together, the OSK and GATA3 analyses indicate that sequence-derived flexibility signatures are detectable at occupied nucleosomal TF-binding sites in vivo. The signal is not explained by simple GC content or dinucleotide composition alone, but its interpretation remains correlative because in vivo occupancy is also shaped by chromatin remodelers, histone modifications, DNA methylation, cooperative TF binding, and local genomic context.

### Flexibility features improve classification performance of TF occupied nucleosomal binding sites across cellular context

Finally, we asked whether the flexibility signal observed for OSK and GATA3 extends to a broader panel of TFs in vivo. We built a classification framework by integrating ChIP-seq occupancy datasets for selected TFs from the ENCODE portal with matched MNase-seq-defined nucleosomal regions from the corresponding cell lines (**Fig. 6a**). Sequence-only models were compared with flexibility-augmented models to test whether DNAflexpy descriptors improve discrimination of occupied TF-binding sites from unbound motif instances present in nucleosomal DNA.

**Figure 6.**
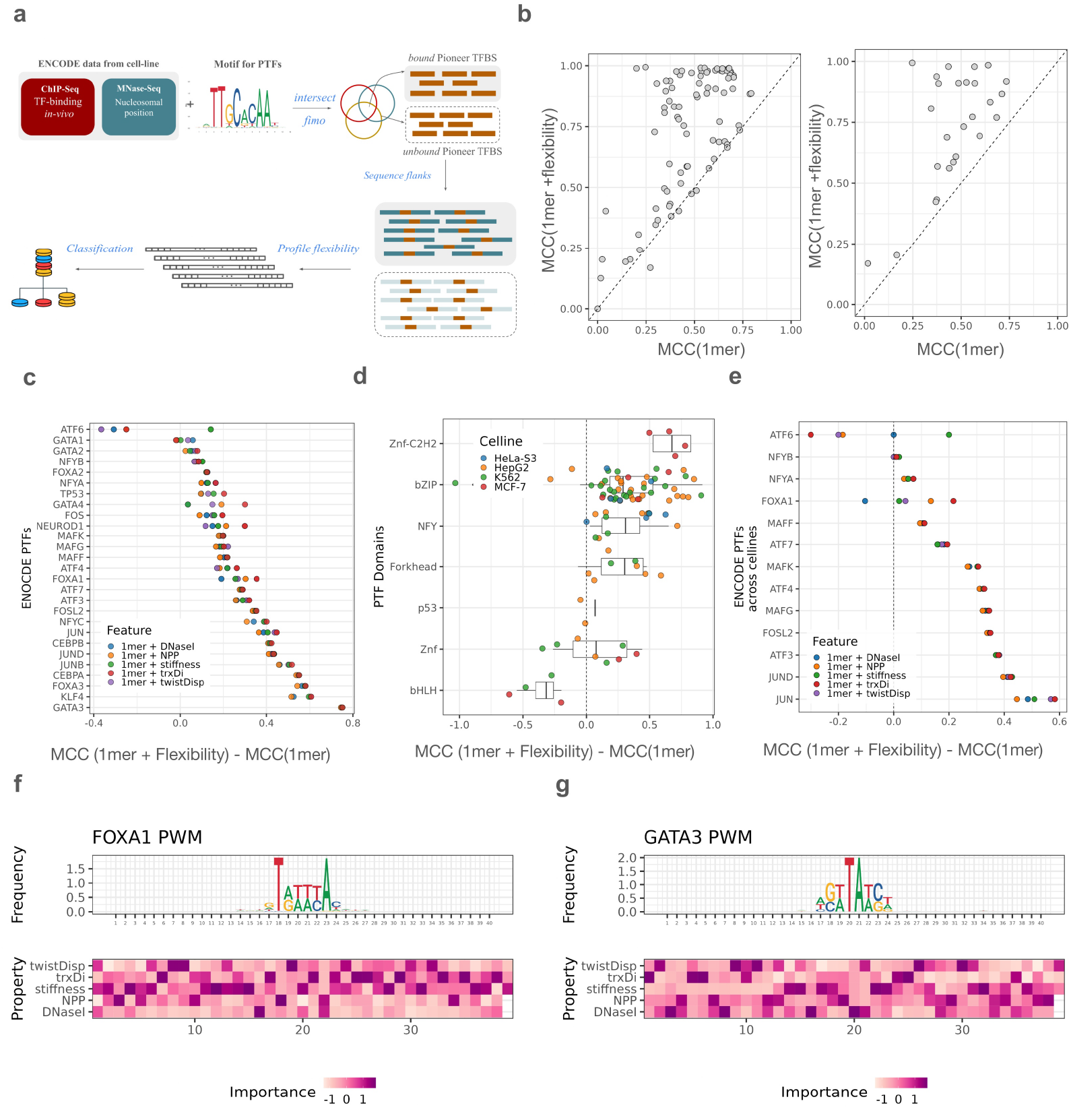
DNA flexibility features enhance machine-learning classification of occupied *in vivo* PTF nucleosomal binding sites. (a) Classification workflow. ENCODE ChIP-seq peaks for 26 pioneer or chromatin-sensitive TFs across multiple cell lines were combined with MNase-seq-defined nucleosomal regions to define occupied nucleosomal positive sites. Matched negatives were unbound motif instances within the same nucleosomal regions and were down-sampled to a 1:1 ratio. XGBoost classifiers used 1-mer one-hot sequence features alone or 1-mer plus DNAflexpy feature sets, and performance was assessed by Matthews correlation coefficient (MCC) under cross-validation. (b) Per-dataset performance for sequence-only models versus flexibility-augmented models. Left: MCC across all ENCODE datasets. Right: median MCC per TF aggregated across cell lines. Points above the diagonal indicate improved classification after flexibility augmentation. (c) ΔMCC = MCC1-mer+feature − MCC1-mer per TF for each DNAflexpy descriptor: DNase I sensitivity, NPP, twist dispersion, trx, and stiffness. Torsional descriptors, especially twist dispersion and trx, produce the largest median improvements. GATA3, KLF4, and FOXA3 show clear positive gains, whereas GATA1, NEUROD1, and ATF6 show limited improvement. (d) Median ΔMCC by DBD family for twist dispersion and trx, with points colored by cell line. bZIP, NFYA, and Forkhead families show the strongest gains in this analysis. (e) Cross-cell-line ΔMCC per TF for individual flexibility features. Models were trained in one cell line and tested on the same TF in another cell line. Torsional and rotational descriptors show the strongest transfer across cellular contexts. (f) Position-specific feature-importance map for the FOXA1 HepG2 model trained with 1-mer + flexibility features using ±20 bp flanks. The top panel shows the sequence logo from positive sites, and the lower panel shows flexibility feature importance by position. FOXA1 weights downstream-flank torsional flexibility most strongly. (g) Equivalent feature-importance map for GATA3. Informative features include upstream torsional descriptors and downstream bending-related descriptors.

Flexibility augmentation improved Matthews correlation coefficient (MCC) across multiple datasets relative to sequence-only models (**Fig. 6b**, left panel). The improvement was clearer when median MCC values were summarized across cell lines for each TF (**Fig. 6b**, right panel), indicating that the signal was not restricted to isolated datasets. Among the flexibility descriptors, torsional features, particularly twist dispersion and twist-roll-X displacement (trx), produced the largest median gains across the 26 TFs analysed. GATA3, KLF4, and FOXA3 showed clear positive ΔMCC with trx features (**Fig. 6c**), whereas GATA1, NEUROD1, and ATF6 showed minimal improvement.

The gains were also structured by DNA-binding domain (DBD) family. Flexibility augmentation was most pronounced for bZIP, NFYA, and Forkhead factors, and bZIP TFs showed improved classification across different cellular contexts, including HepG2, K562, MCF-7, and HeLa-S3 (**Fig. 6d**). We next tested cross-cell model performance by training models for a given TF in one cell line and evaluating classification performance in another. Flexibility-augmented models improved performance for some TFs, although the effect was not uniform across cellular contexts (**Fig. 6e**). Thus, flexibility features might represent a transferable component of binding-site discrimination for selected factors, but this contribution remains TF- and context-dependent.

Position-specific feature analyses further showed that the flexibility signal was spatially organized around motifs. For FOXA1, torsional flexibility in one motif flank was among the most informative features, whereas GATA3 showed contributions from both upstream torsional and downstream bending-related descriptors (**Fig. 6f,g**). These patterns suggest that different DBDs may use distinct motif-proximal structural contexts during nucleosomal engagement.

Overall, this classification analysis extends the biochemical modelling and in vivo case-study results to a broader TF panel. Flexibility-augmented models modestly but reproducibly improved discrimination of occupied nucleosomal binding sites for selected TFs and cell types. Torsional descriptors emerged as the most informative features within the current dataset and classification framework, supporting the view that DNAflexpy flexibility profiles capture one layer of functional site-selection information among multiple chromatin and sequence determinants.

## Discussion

Precise recognition of cis-regulatory elements by transcription factors (TFs), together with their regulated occlusion within chromatin, underlies eukaryotic gene regulation [74]. In a chromatinized genome, a motif is not read in isolation. Nucleosomes present DNA as a sharply bent, rotationally phased substrate that is periodically contacted by the histone octamer [81]. Thus, a cognate motif is interpreted within a mechanically constrained sequence context [17,45]. Motif position, rotational orientation [14], architecture of DNA-binding-domain (DBD) of TFs [77], and histone modifications [8] all influence TF engagement with nucleosomal DNA. Studies on TF binding specificity have further shown that sequence models can only partially explain their binding preferences and affinity landscape [7]. DNA shape and conformational flexibility of flanking sequences can modulate affinity even at positions not directly contacted by the protein [37,38]. Several TF-bound nucleosome structures suggest that such structural information remains relevant in nucleosomal DNA [19–22]. However, a systematic test of how DNA flexibility is relevant for TF binding in nucleosome has been limited [46,82–84], especially across TFs with different DNA binding domains, and distinct nucleosomal context [15]. This gap is also relevant as local DNA structure and genomic features are also known to modulate the TF binding specificity [7,74,85].

Here, we address this gap by showing that sequence-derived flexibility descriptors provide an interpretable layer for accurately predicting TF-nucleosome affinity across a broad TF panel. The framework identifies deformation-sensitive positions, reveals family- and template-dependent structural signatures, and remains detectable at occupied nucleosomal sites in vivo. These observations are consistent with a model in which sequence-encoded mechanics may tune the energetic cost of local deformation during TF-nucleosome recognition [40]. Productive engagement may require motif exposure, sterically compatible DBD binding, local bending or groove remodeling, and stabilization of the bound state against histone-DNA constraints [86]. The descriptors do not measure these events directly. Rather, they may report sequence contexts that are more compatible with such deformations.

The predictive gain can be interpreted through the deformation modes encoded by the descriptors. DNase I sensitivity reports local bendability, particularly the tendency of trinucleotide steps to bend toward the major groove [38,87]. In nucleosomal TF binding, this may mark motif-proximal sequences that can accommodate bending strain during exposure, local distortion, or induced fit. NPP provides a related but distinct signal [30,88]. It reflects trinucleotide preferences associated with rotational orientation on the histone surface [89]. It may therefore capture whether local sequence favours a groove-facing register compatible with wrapping and TF access. Twist dispersion and trx add torsional information, including coupled changes in twist, roll, and base-step displacement [31]. These properties are relevant since many TFs require local twist adjustment, groove presentation, or base-step reorientation in addition to bending. Stiffness provides a complementary term by reporting resistance to stretching or compression [84,90]. Together, these descriptors offer a compact representation of deformability, phasing, torsional adjustment [91], and resistance to distortion [38,61,92,93].

This interpretation also explains why the gain is not uniform across TF families. A descriptor is expected to improve prediction only when the deformation mode it represents is relevant to the binding strategy of that TF. SOX/HMG factors are likely to be more sensitive to deformability because their binding involves sharp bending and minor-groove remodeling [69,70]. Other families, including GATA, bZIP, Homeodomain, CUT, RHD, Forkhead, and zinc-finger factors, are known to employ distinct mechanisms of nucleosomal engagement [14], and may therefore utilize the flexibility information like groove presentation, torsional adjustment to a differing extents in the nucleosomal DNA. The same feature may therefore carry different predictive weight across DBD families. This is consistent with the observation that structural preferences are not universal, even among TFs with broadly similar DNA-binding folds [38].

Flexibility descriptors are sequence-derived and should not be treated as physical measurements independent of nucleotide composition. Rather, they reorganize sequence information into a mechanistically interpretable feature space [61]. Their largest gains were observed over mononucleotide baselines, with smaller residual gains over trinucleotide baselines. The comparisons highlight the plausible importance of DNA flexibility as a efficient encoding of sequence dependent structural information not as a ideal replacement of sequence features. The comparison with ShapeFL, derived using Deep DNAshape, supports this interpretation [63]. Deep DNAshape predicts sequence-dependent fluctuations in shape parameters from extended flanking context. DNAflexpy encodes empirical deformation tendencies from DNase I cleavage, nucleosome-positioning preferences, and base-step geometry. Despite these distinct origins, the two representations recovered overlapping TF-nucleosome datasets. Their partial non-redundancy may reflect complementary sensitivities: local step-level axial, torsional, and stretching tendencies for DNAflexpy, and learned shape variability over broader flanks for ShapeFL. The convergence argues against a feature-specific artifact and points to a shared structural signal in nucleosomal sequence context. Other models, including DNAcycP, and pyDNAxEPBD, were not benchmarked here [94–97]. Direct comparison with these models may further clarify whether they provide overlapping or complementary information.

Beyond these different structural representations, the flexibility signal also appeared to depend on the nucleosomal template itself. In the PIONEAR-seq datasets, flexibility-associated gains differed between the W601 and genomic templates, and cross-template prediction performance varied among TFs. For a given TF, position-resolved flexibility footprints appear to emerge from the combined influence of motif sequence, flanking context, and nucleosomal template [15,46,64]. PAX7 and GATA4 retained footprints that remained largely centred on similar motif-proximal steps across the three genomic templates (*Fig. S5a,b*). In contrast, FOXA1 and PU.1 showed weaker footprints, with stronger divergence on W601 than on the genomic templates (*Fig. S5c,d*). These observations suggest that the same cognate motif can carry different structural readout depending on its nucleosomal positioning sequence. Thus, the nucleosomal template may impose additional structural constraints that alter which motif proximal steps are most informative for deformation compatible TF engagement.

The SOX-nucleosomal binding case is where the footprints connect most directly to a structural and dynamic mechanism, although the evidence stays correlative. Regression-derived flexibility footprints localized to the SOX motif core and flanks, matching the region that undergoes minor-groove remodeling in the SOX11-DNA structure and separates high-from low-affinity sites by MGW [19]. MD simulations linked this footprint to a dynamic binding pathway [64]. At SHL2, the SOX-unbound nucleosome showed elevated MGW fluctuation around the motif, suggesting a broader pre-binding ensemble. Upon SOX binding, the motif adopted a widened and less-fluctuating minor groove, consistent with bound-state stabilization. This pattern is compatible with coupled conformational selection and induced fit. Pre-binding flexibility may increase sampling of SOX-compatible states, while binding may further remodel and stabilize the DNA. Because the regression model was trained on neither structural nor MD data, this agreement gives the mechanistic support beyond prediction method.

These findings may facilitate the interpretation of regulatory variants. Many noncoding variants preserve the core motif while altering flanking mechanics, phasing, bendability, or groove presentation. Such variants could change motif usability in chromatin without changing motif identity. Deep-learning models that can predict variant effect model statistical sequence patterns, but flexibility descriptors may help explain how variants alter local structural compatibility within nucleosomal DNA [98,99]. The principal limitation remains that flexibility descriptors are sequence-derived. Their predictive value does not establish flexibility as a causal variable separable from sequence composition. Trinucleotide baselines, ShapeFL comparisons, dinucleotide-preserving controls, and cross-template tests mitigate this concern to an extent, but sequence, shape, curvature, periodicity, and mechanics remain intertwined.

The most direct next step is experimental perturbation. Designed nucleosomal templates that preserve motif identity while altering flanking flexibility, rotational phase, or bendability would test whether predicted footprints can be shifted, weakened, or transferred between contexts. Longer replicated MD simulations could help resolve when a sequence is compatible with conformational selection, induced fit, or post-binding stabilization. To summarise, sequence-encoded DNA flexibility is not a replacement for sequence models, DNA shape, or chromatin accessibility. It is an interpretable layer of indirect readout that may help explain why only a subset of motif instances becomes mechanically usable within nucleosomal DNA.

## Conclusions

Our analyses establish DNA conformational flexibility as a quantifiable, sequence-encoded layer of the transcription factor–nucleosome recognition landscape. Across NCAP-SELEX and PIONEAR-seq, flexibility-augmented regression models systematically improve prediction of TF–nucleosome binding affinity over mononucleotide baselines and retain residual gains over trinucleotide composition, with effect sizes that differ reproducibly across DNA-binding domain families. Position-resolved flexibility footprints derived from the same models converge on positions of direct structural deformation in the SOX11–DNA cocrystal and on register-specific minor-groove dynamics observed in all-atom MD trajectories, supporting a mechanistic, rather than purely empirical, interpretation of the predicted signal. In chromatin, flexibility signatures distinguish nucleosomal from non-nucleosomal OSK binding sites and pioneered from non-pioneered GATA3 regions, and improve classification of occupied PTF binding sites across multiple ENCODE cell lines and PTF families. These associations remain correlative and do not establish flexibility as a sole causal determinant of pioneering activity; however, they position sequence-derived DNA mechanics as a compact, interpretable representation of the indirect-readout layer governing pioneer transcription factor engagement with nucleosomal DNA.

## Methods

### NCAP-SELEX data processing

Raw sequencing data (FASTQ format) from NCAP-SELEX, HT-SELEX, and Nucleosome-SELEX experiments were obtained from the European Nucleotide Archive (ENA; accession PRJEB22684) [14]. Quality control (QC) and preprocessing were performed using fastp v1.3.1[100]: adapter sequences were trimmed, reads containing ambiguous nucleotides (N) were removed, PCR duplicates were discarded, and forward and reverse reads were merged. Only merged reads of exactly 101 bp, corresponding to the randomized region of the SELEX library template, were retained; reads of other lengths were discarded.

*k*-mer binding affinities were computed from quality-controlled NCAP-SELEX reads with the R SELEX package v1.44.0 [101]. For each TF, cycle 4 was compared with the cycle 0 input library to estimate sequence specific enrichment. The variable region length was inferred from each library’s read length. A Markov background model was fit to cycle 0 with DIVISION cross-validation and automatic order selection. This step corrected higher-order sequence composition biases. Final *k*-mer affinities were calculated as enrichment over the Markov expectation. A minimum count filter equal to *k* was applied for robust estimation.

To capture motif-proximal context, we used iterative flanking expansion rather than a fixed *k*-mer length. Starting from each TF’s IUPAC-encoded reference motif [14], we added one flanking base on each side per iteration. The initial length was *k* ≈ 10. Affinity tables were generated at each flank length. Expansion stopped when fewer than 1,000 *k*-mers had measurable affinity. This procedure produced affinity tables with increasing flanking-context resolution.

Affinity tables were filtered to retain motif-related *k*-mers. For each TF and flank length, the IUPAC-encoded reference motif [14] was padded with flanking N positions. The allowed mismatch threshold was computed from the effective consensus length. Each position contributed 1/*d*_*i*_, where *d*_*i*_ is the number of nucleotides encoded at position *i*. The threshold was *m* = max(0, ⌊(*L*_eff_ − 4)/2⌋ + 1), where *L*_eff_ = ∑_*i*_ 1/*d*_*i*_. We built a trie data structure from affinity-table *k*-mers for prefix-based pruning during candidate enumeration. Candidate sequences were generated by depth-first search over the expanded reference. Prefix filtering avoided branches absent from the affinity table. Only *k*-mers present in both the candidate set and affinity table were retained.

Filtered *k*-mer sets were aligned using the top-down-crawl (TDC) algorithm (https://github.com/bhcooper/TopDownCrawl [102]. Briefly, TDC first merges each k-mer with its reverse complement by averaging their affinity scores, then seeds the alignment from the highest-affinity k-mer. Remaining k-mers are assigned positional shifts relative to the seed through three operations applied iteratively in affinity-ranked order: (i) single-nucleotide variant matching (identifying k-mers differing by one substitution), (ii) 1 bp sliding-window overlap (left or right extension), and (iii) 2 bp sliding-window overlap for more distant relationships. K-mers that cannot be connected to the aligned set are reported separately as unaligned. TDC was applied separately for each flank length (0, 1, and 2 flanking positions), processing only tables exceeding 1,000 k-mers to ensure sufficient data for robust alignment. From the aligned k-mer sets, affinity-weighted position weight matrices (PWMs) were generated as information-content sequence logos [103], trimming edge positions with sparse coverage (up to 2 positions per side). Aligned k-mer tables with their affinity value were directly used for downstream analysis.

### PIONEAR-seq data processing and modelling

PIONEAR-seq data were obtained from GEO accession GSE293487 [15]. The assay tiles a 20-bp randomized window (*N*_20_) across four 147-bp nucleosomal templates: the synthetic Widom 601 positioning sequence (W601) and three genomic positioning sequences, ALB1, CX3, and NRCAM. Libraries were assembled into reconstituted nucleosome core particles and subjected to four rounds of affinity selection with full-length human transcription factors.

We used the authors’ processed per-round *k*-mer enrichment tables directly. Each table reports a normalized enrichment value (`Increase`) for every observed *k*-mer relative to the initial unselected library. Both nucleosome-associated (NCP) and free-DNA control (DNA) tables were retrieved. Data for 11 TFs across four templates were retained, yielding 44 Round 4 NCP TF-template datasets (11 TFs × 4 templates). Biological replicates were available for FOXA1, GATA1, GATA4, OCT4, PAX7, and RUNX2.

For each TF-template-replicate combination, nucleosome-specific *k*-mer affinity was defined as:

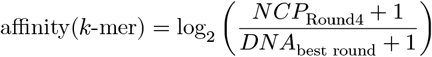

where *NCP*_Round4_ is the `Increase` value from the Round 4 NCP pulldown table, and *DNA*_best round_ is the `Increase` value from the matched free-DNA control table. A pseudocount of 1 was added to both terms to stabilize low-enrichment values. For the DNA denominator, Round 1 was used when available because it is the least-enriched free-DNA control. If Round 1 was unavailable for a given TF-template-replicate combination, Round 3 was used. The *k*-mer length followed the assay design: 10-mer for OCT4; 9-mer for FOXA1, KLF4, PU.1, and ZNF143; and 8-mer for the remaining TFs.

Each TF-template FASTA was modelled using the same nested L2-regularized ridge regression framework used for NCAP-SELEX. We used 10-fold outer and 5-fold inner nested cross-validation. The ridge penalty parameter was selected in the inner loop from 500 log-spaced values between 10^−3^ and 10^7^. Target affinity values were min-max scaled within each TF-template dataset before fitting. Model performance was summarized as mean cross-validated *R*^2^ across outer folds, and Cohen’s *f* ^2^ was computed as described for NCAP-SELEX.

For cross-replicate concordance, models were trained on one biological replicate and evaluated on the other replicate, in both directions. Spearman rank correlation between predicted and observed affinities on the held-out replicate was used as the primary metric because it is insensitive to differences in absolute scale. For cross-template generalization, models were trained on one nucleosomal template and evaluated on each of the remaining three templates for the same TF. This produced 12 directed train-to-test template pairs per TF and followed the same ridge-regression protocol.

### DNAflexpy software and DNA flexibility descriptors

DNA flexibility descriptors were computed with DNAflexpy, an open-source Python package developed to convert DNA sequences into position-resolved flexibility profiles (https://github.com/UpalabdhaD/DNAflexpy). DNAflexpy maps each overlapping di- or trinucleotide in a sequence to a value from an internal lookup table and returns a numeric profile that can be used either alone or concatenated with sequence encodings for regression and classification.

Five descriptors were used throughout the study. DNase I bendability is a trinucleotide scale derived from sequence-dependent DNase I cleavage preferences and was used as an empirical proxy for local bendability [87]. Nucleosome positioning preference (NPP) is also trinucleotide-based and captures sequence periodicity associated with nucleosome positioning and rotational bending preferences [89]. Twist dispersion is a dinucleotide-step descriptor that summarizes variability in twist angles observed across DNA-protein structural contexts and was used as a torsional-flexibility measure. The twist-roll-X displacement descriptor (trx) is a dinucleotide-step composite that combines twist, roll, and X-displacement information, capturing coupled torsional, bending, and translational deformation [31]. Stiffness modulus was used as a stretching or compressibility-related descriptor derived from elastic parameters of DNA duplex deformation. DNase I sensitivity and NPP therefore use 64 unique trinucleotide lookup tables, whereas twist dispersion, trx, and stiffness use 16 unique dinucleotide lookup tables. The source scales have different native units and ranges; after feature construction, all descriptors were treated as normalized, unitless model inputs. DNase I sensitivity, NPP, twist dispersion, and trx were interpreted as flexibility or deformation-propensity descriptors, whereas stiffness was retained as an elastic-resistance descriptor whose contribution was learned by the model.

Unless otherwise stated, flexibility profiles were computed at single-base resolution using DNAflexpy with window size 0. For each sequence, descriptor values were min-max scaled. To capture local dependencies between adjacent positions, lag-1 interaction terms were generated as the product of consecutive position values for each descriptor. The original flexibility profiles and lag-1 terms were concatenated and rescaled before being combined with the *k*-mer (1-mer, 3-mer) encodings.

### DeepDNAshape shape-fluctuation feature generation

Deep DNAshape shape-fluctuation features were generated with the publicly released Deep DNAshape v0.1 [63] (https://github.com/JinsenLi/deepDNAshape). Current analysis used shape fluctuation outputs (‘-FL’), which estimate sequence-dependent fluctuation of DNA shape parameters. ShapeFL profiles used here include minor groove width fluctuation (MGW-FL), propeller twist fluctuation (ProT-FL), helix twist fluctuation (HelT-FL), roll fluctuation (Roll-FL), and slide fluctuation (Slide-FL).

For each aligned binding-site sequence, a FASTA record was supplied to Deep DNAshape and per-position fluctuation values were returned for each requested layer. We evaluated layers L0, L2, and L5, corresponding to predictions made with 0-, 2-, and 5-nt extended flanking sequence context around each predicted base or base-pair step. ShapeFL tracks were obtained from the same TDC-derived binding-site used for DNAflexpy features.

### Multiple linear regression

To relate binding affinity with DNA sequence and flexibility features, we employed *L*_2_-regularized (Ridge) multiple linear regression (L2-MLR) in a nested cross-validation framework, following the general approach of concatenating sequence encoding with flexibility features [61,62]. For an affinity vector *y* with *n* training observations and *p* encoded features, the regression model is

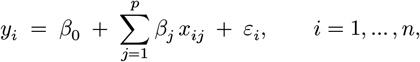

where *β*_0_ is the intercept, *β*_*j*_ are regression coefficients on the encoded sequence and flexibility features *x*_*ij*_, and *ε*_*i*_ is the residual error. To prevent overfitting in this high-dimensional regime, the coefficient vector *β* is obtained by minimizing the *L*_2_-penalized loss

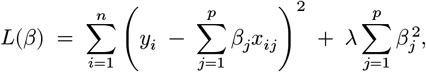

where *λ* ≥ 0 controls the strength of *L*_2_ regularization and is selected by inner cross-validation (see below).

DNA sequences were represented using one-hot encoding at multiple *k*-mer resolutions. For a sequence of length *L* and a given *k*, each of the *L* − *k* + 1 overlapping *k*-mer windows was encoded as a binary vector of length 4^*k*^, yielding a feature vector of dimension (*L*−*k*+1) ×4^*k*^. We evaluated models at *k* = 1 (1-mer, 4 features per position), *k* = 2 (2-mer, 16 features per position), and *k* = 3 (3-mer, 64 features per position), as well as a combined encoding concatenating all three resolutions (1-mer + 2-mer + 3-mer). Positions containing gap characters introduced during TDC alignment were encoded as all-zero vectors.

Five DNA flexibility descriptors were computed for each sequence using the DNAflexpy package: DNase I sensitivity [87], nucleosome positioning preference (NPP) [89], twist dispersion, the twist-roll-X-displacement (trx) composite, and stiffness modulus. Each feature was computed at single-base resolution (window size = 0) and scaled row-wise to [0, 1] using min-max normalization. To capture local dependencies between adjacent positions, lag-1 interaction terms (the element-wise product of each pair of consecutive positions) were appended to each feature vector, and the combined feature matrix was rescaled row-wise.

We evaluated seven model configurations to assess the contributions of sequence encoding and flexibility features: 1-mer only; 2-mer only; 3-mer only; combined 1-mer + 2-mer + 3-mer; flexibility only (without sequence encoding); 1-mer + flexibility; and 3-mer + flexibility. This design allows direct comparison of whether flexibility features provide predictive information beyond what is captured by *k*-mer sequence encoding alone, including at the 3-mer level which inherently captures nearest-neighbor dinucleotide effects [62].

Target affinities were min-max scaled to [0, 1] for each dataset. Each dataset (TF × flank combination) was modeled independently, as relative affinities may not be comparable across different SELEX experiments. We used Ridge regression (scikit-learn `Ridge`) within a nested cross-validation design: the outer loop comprised 10 folds, and within each outer training set, the regularization parameter *λ* (termed `alpha` in scikit-learn) was selected by 5-fold inner cross-validation over 500 candidate values spaced log-uniformly from 10^−3^ to 10^7^. The inner fold achieving the highest mean *R*^2^ determined the optimal *λ*, which was then used to refit the model on the full outer training set and predict held-out outer test fold affinities. Model performance was assessed using the coefficient of determination (*R*^2^), mean squared error (MSE), and mean absolute error (MAE) across the 10 outer folds.

### Quantifying nonredundant model performance using Cohen’s *f* ^2^

To quantify the nonredundant contribution of DNA flexibility descriptors beyond sequence-only baselines, we computed Cohen’s *f* ^2^ effect sizes from mean cross-validated *R*^2^ values, following standard hierarchical regression conventions [61]. For a flexibility-augmented model (e.g., 1-mer + flexibility) and a corresponding nested sequence-only baseline (e.g., 1-mer), *f* ^2^ is defined as

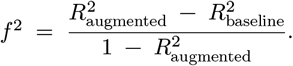

This quantity expresses the proportion of variance uniquely attributable to the augmentation relative to the residual variance left unexplained by the augmented model. Per-dataset *f* ^2^ values were computed for each TF, then summarized at the DBD-family level. Following standard conventions, *f* ^2^ ≥ 0.02, ≥ 0.15, and ≥ 0.35 were interpreted as small, medium, and large effect sizes, respectively. The analysis was applied both over the 1-mer baseline, to quantify the marginal contribution of flexibility beyond mononucleotide identity, and over the 3-mer baseline, to assess the incremental validity of flexibility beyond trinucleotide composition. The same framework was used to compare DNAflexpy with Deep DNAshape-derived shape-fluctuation (ShapeFL) features [63] at layers L0, L2, and L5 over both 1-mer and 3-mer baselines.

### Position-specific flexibility contributions

To resolve the contribution of DNA flexibility at individual nucleotide positions within the binding site, we adapted a position-specific decomposition strategy previously developed for shape-readout analysis on free DNA [36,61]. For each TF, we first fit a baseline 1-mer ridge regression model and recorded its cross-validated coefficient of determination, 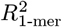. We then iteratively augmented the feature set by appending the five flexibility scores (DNase I sensitivity, NPP, twist dispersion, trx, and stiffness) at a single nucleotide position *i*, refit the model under the same nested cross-validation protocol, and obtained the new cross-validated performance 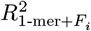. No interaction terms were included in this position-resolved decomposition.

The performance gain attributable to flexibility at position *i* is

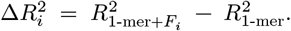

To express the gain on a common scale across TFs with different baseline *R*^2^, we computed the relative contribution at position *i* as

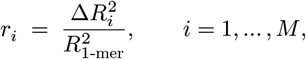

where *M* is the binding site length. A positive *r*_*i*_ indicates that flexibility at nucleotide *i* improves affinity prediction beyond mononucleotide identity, defining a position-specific flexibility readout. Plotting the *r*_*i*_ profile across positions for each TF yields a TF-specific flexibility footprint; aggregating these footprints across DBD families produced heatmaps highlighting recurrent positions of bendability (DNase I, NPP), torsional (twist dispersion, trx), and stretching (stiffness) flexibility contribution to nucleosomal binding specificity. The same procedure was applied to PIONEAR-seq datasets to derive template-resolved flexibility footprints, and the resulting positional contribution maps were compared with the SOX11-DNA complex structure (PDB: 6T78) and all-atom MD trajectories of SOX-bound and SOX-free nucleosomes at SHL0, SHL2, and SHL4.

### Analysis of MD simulations of SOX-bound nucleosomes

All-atom MD trajectories of SOX-bound and SOX-unbound nucleosomes were taken from the public dataset of Ozden *et al*. [64]. As reported in the source dataset, simulations used distinct force fields for protein and DNA, Amber ff14SB for protein, parmbsc1 for DNA, TIP3P water. Additionally, system were solvated in 0.15 M NaCl at 310 K, production runs were performed with a 2 fs integration step, and snapshots were written every 0.5 ns.

We reanalysed eight trajectories of total 5 µs: two replicas of 1000 ns for the SOX-unbound mutated nucleosome on the W601 sequence (W601-SHL024), in which SOX11 motifs were inserted at SHL 0, 2, and 4, and two replicas of 500 ns for SOX11 bound at each of the same three SHL positions (6 × 500 ns).

Equilibration frames were excluded prior to analysis, following Ozden *et al*.. Frames before 350 ns were discarded for W601-SHL024 (SOX unbound nucleosomal simulations), and 200 ns for SOX-bound systems. For each remaining frame, we extracted a 20-bp DNA window centered on the inserted SOX11 motif, comprising the 12-bp insertion (5^′^-GGACAATGGAGG-3^′^) flanked by 4 bp on either side. DNA groove widths and base-step parameters were computed across this window with X3DNA v2.4.8-2023nov10 [104]. Per-position profiles for DNA shape parameters (MGW, Slide, Roll, Twist, Tilt, Shift, Rise, helical rise, and helical twist) were obtained by averaging values over the frames from the post-equilibration time.

Local DNA conformational flexibility was estimated from per-position shape fluctuations. For each system (bound/unbound), superhelical location (SHL 0, 2, 4), replicate, and base position in the 20-bp window, we computed the within-replica standard deviation across post-equilibration frames. Per-replica values were then combined into per-position MGW fluctuation (MGW-FL) profiles using the pooled standard-deviation formula,

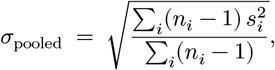

where *s*_*i*_ is the within-replica standard deviation and *n*_*i*_ is the number of post-equilibration frames in replica *i*.

Time-resolved MGW analyses used the same per-frame X3DNA groove-width values. The core step (the central `CA/TG` step of `ACAA`) and the downstream step were tracked over time, with LOESS smoothing applied for visualization. Time-resolved MGW heatmaps at each SHL were generated by averaging frames into 5 ns bins, and ΔMGW-FL heatmaps were computed as the difference between bound and unbound pooled standard deviations over 50 ns sliding windows sampled every 10 ns.

Shape and shape-fluctuation distributions for SOX-bound versus unbound systems were compared using two-sided Mann–Whitney U tests on per-frame observations, with Benjamini–Hochberg correction applied for each panel.

### Genomic analysis of OSK binding sites from ChIP-seq and MNase-seq data

For the OSK analysis, OCT4, SOX2, and KLF4 ChIP-seq binding sites in IMR90 cells were taken from Soufi et al. [16] (GEO accession GSE36570). To assign each site a local nucleosome state, we used matched MNase-seq from Kelly et al. 2012 (GSE21823) [105]. BAM files were pooled with SAMtools [106] and converted to RPGC (Reads Per Genomic Content) normalized bigWig tracks with deepTools (v3.5.5) `bamCoverage` [107], binned at 10 bp. Sites were then split on mean MNase signal in a 200-bp window around each peak summit; signal below 1 we called nucleosome-depleted, the rest referred as nucleosome enriched. Heatmaps used a ±1-kb window around each summit. Before partitioning, the OSK datasets contained 22405 OCT4, 39397 SOX2, and 27988 KLF4 sites.

Sequences ±500 bp around each peak summit were extracted with BEDTools [108] using hg18 coordinates and lifted to hg38 first where needed. Flexibility profiles were computed using DNAflexpy package as described above. GC content between nucleosomal and non-nucleosomal sites was compared with Wilcoxon rank-sum tests; we used Cliff’s *δ* value to obtain the effect size. To check the local nucleotide composition of the flexibility profiles, sequences were shuffled with preserving dinucleotides with biasaway v3.3.0 [109], and flexibility profiles recomputed on the shuffled sequences.

For the OCT4 occupancy analysis, nucleosomal OCT4 sites were ranked into deciles by either mean DNase I sensitivity or twist dispersion across the peak-centered window. Kruskal–Wallis tests then compared ChIP-seq signal across deciles, and the top-versus-bottom contrast was reported with Cohen’s d and 95 CIs. Partial Spearman correlations between ChIP-seq signal and flexibility were computed with ppcor (`pcor.test`), controlling for GC content.

To ask whether sequence-derived structural features predict local nucleosome occupancy, XGBoost regression models were trained on the continuous MNase-seq signal in the 200-bp summit-centered window. Five feature sets were compared: 1-mer sequence, 3-mer sequence, flexibility-only, ShapeFL-only, and the combined structural encoding. Performance was assessed by 10-fold stratified cross-validation, with binary nucleosomal labels held back for the post hoc ROC and precision–recall analysis only (XGBClassifier, xgboost; `n_estimators=300, max_depth=6, learning_rate=0.1, subsample=0.8, colsample_bytree=0.8, eval_metric=‘logloss’, random_state=42; StratifiedKFold(shuffle=True, random_state=42)`).

### Flexibility profiling of GATA3 binding sites from ATAC-seq and ChIP-seq data

For GATA3, we used ChIP-seq peaks and ATAC-seq reads from MDA-MB-231 cells with and without ectopic GATA3 expression (GEO GSE72141) [75]. Replicate FASTQs were trimmed for adapters and quality with fastp [100], aligned to hg19 with Bowtie2 (`--very-sensitive --no-mixed --no-discordant -X 5000`) [110], then sorted and indexed with SAMtools v1.3.1[106]. featureCounts v2.0.8 [111] gave read counts in ±100-bp windows around each GATA3 ChIP-seq peak center, and DESeq2 v1.46.0 [112] tested differential accessibility between the GATA3-expressing and wild-type conditions under a two-group design.

Each GATA3-bound region was placed in one of three accessibility classes based on its response to GATA3 induction: *pioneered* sites passed FDR < 0.05 with log2 fold-change > 2; constitutively *open* sites had FDR > 0.05 and mean normalized ATAC-seq counts above a threshold of 20 in both conditions; and closed or non-pioneered sites had FDR > 0.05 with mean normalized counts below 10 in both. Coordinates were lifted to hg38, and ±500 bp around each peak or motif center was used for flexibility profiling using DNAflexpy. Motif occurrences themselves were located with FIMO at p < 0.001 [113] using JASPAR matrix for GATA3 (MA0037.3) [114,115]. Per-position flexibility profiles were then averaged across sites within each class. For rotational positioning, WW/SS dinucleotide scores were computed from the Cui–Zhurkin model [79] over the same ±500-bp motif-centered windows using a custom python script: with a 147-bp sliding window at step 1. The output is an 854-position profile spanning −427 to +426; scores were normalized per sequence, without any smoothing.

### Genome-wide classification of TF binding sites in nucleosomal regions with XGBoost

For the genome-wide classification analysis, we curated ChIP-seq peak sets for 26 pioneer or chromatin-sensitive transcription factors across HepG2, K562, MCF-7, HeLa-S3, and a handful of other ENCODE cell contexts. IDR-thresholded ChIP-seq peaks and matched MNase-seq peak files were pulled from the ENCODE portal. For each TF, JASPAR motif profiles [114,115] were scanned across hg38 with FIMO [113] (MEME Suite v5.5.7; p < 0.0001).

Positives were motif instances that fell within both an ENCODE ChIP-seq peak for the corresponding TF and an MNase-seq nucleosomal region in the same cell line. Negatives were drawn from motif instances in the same nucleosomal regions that did not intersect the corresponding TF peak set, then randomly down-sampled to a 1:1 ratio with positives. Sequences around each motif instance were extracted from hg38 with BEDTools [108] using 10- or 20-bp flanks; the main feature-importance maps used the 20-bp setting.

Each sequence was encoded two ways: as 1-mer one-hot vectors and as DNAflexpy-derived flexibility profiles. We compared a 1-mer baseline against 1-mer plus each individual flexibility descriptor (DNase I sensitivity, NPP, twist dispersion, trx, or stiffness), and against a combined model concatenating all five with the 1-mer encoding. XGBoost classifiers were trained through the scikit-learn API. Hyperparameters were first tuned with GridSearchCV on a pooled HepG2 training set using 1-mer + DNase I features; the pool was split 80:20 into train and validation, with 10-fold stratified CV inside the training half (XGBClassifier, xgboost; grid: `n_estimators {50,100,200}, max_depth {3,6,9}, learning_rate {0.01,0.1,0.2}, subsample {0.8,0.9,1.0}, colsample_bytree {0.8,0.9,1.0}`; best fit: `n_estimators=200, max_depth=9, learning_rate=0.1, subsample=0.9, colsample_bytree=0.8, eval_metric=‘logloss’, random_state=42; reg_alpha` and `reg_lambda` left at XGBoost defaults).

The tuned configuration was then applied independently to each TF and cell-line combination under 5-fold stratified cross-validation. We tracked the full panel of metrics: Matthews correlation coefficient (MCC), accuracy, precision, recall, F1, ROC AUC, and precision–recall AUC. For the main comparisons we report MCC and ΔMCC of each flexibility-augmented model relative to the 1-mer baseline. For cross-cell prediction, a model trained on one cell line for a given TF was applied to held-out examples for the same TF in a different cell line, evaluated with the same metrics. ΔMCC came from subtracting the sequence-only baseline from each flexibility-augmented model.

For position-specific interpretation, feature importance was computed from XGBoost models trained on the HepG2 FOXA1 and GATA3 datasets with 20-bp flanking windows. Importance scores for sequence and flexibility features were then aggregated by nucleotide position and descriptor type across folds.

## Declarations

### Ethics approval and consent to participate

Not applicable. This study did not involve human participants, animal subjects, or identifiable patient data. All analyses were performed on publicly available datasets and on molecular dynamics trajectories released by their original authors.

### Consent for publication

Not applicable.

### Availability of data and materials

All datasets analysed in this study are publicly available. A consolidated list of accession identifiers and access URLs is provided below; further details are also recorded in the Methods.

**In vitro TF–nucleosome binding (NCAP-SELEX, HT-SELEX, Nucleosome-SELEX):** European Nucleotide Archive (ENA) project PRJEB22684 [14], available at https://www.ebi.ac.uk/ena/browser/view/PRJEB22684.

**PIONEAR-seq:** Raw and processed data from [15] are available at the Gene Expression Omnibus (GEO) under accession GSE293508 (https://www.ncbi.nlm.nih.gov/geo/query/acc.cgi?acc=GSE293508).

**OSK ChIP-seq and MNase-seq (IMR90 cells):** [16], GEO accession GSE36570 (https://www.ncbi.nlm.nih.gov/geo/query/acc.cgi?acc=GSE36570).

**GATA3 ChIP-seq and ATAC-seq (MDA-MB-231 cells):** [75], GEO accession GSE72141 (https://www.ncbi.nlm.nih.gov/geo/query/acc.cgi?acc=GSE72141).

**ENCODE ChIP-seq and DNase-seq:** Datasets for 26 pioneer transcription factors across HepG2, K562, MCF-7, HeLa-S3, and additional ENCODE cell lines were obtained from the EN-CODE portal (https://www.encodeproject.org/).

**Structural data:** The SOX11–DNA cocrystal structure was retrieved from the Protein Data Bank under accession PDB: 6T78.

**Molecular dynamics trajectories:** All-atom MD trajectories of SOX-bound and SOX-free nucleosomes at SHL0, SHL2, and SHL4 (mutated W601 templates) were obtained from the public repository associated with [64].

**DNA flexibility descriptors:** Five flexibility scales (DNase I sensitivity, NPP, twist dispersion, trx, and stiffness) were computed using the open-source Python package

**DNAflexpy**, available at https://github.com/UpalabdhaD/DNAflexpy.

**DNA shape-fluctuation features:** Deep DNAshape shape-fluctuation features (ShapeFL layers L0, L2, and L5) were generated using the publicly released Deep DNAshape software [63], available at https://github.com/JinsenLi/deepDNAshape.

**Analysis code:** All custom scripts used for data preprocessing and analysis are deposited in the github repository with instructions, avaialble at https://github.com/UpalabdhaD/DNAflexibility_in_TF-NucleosomalDNA_MS.git

### Authors’ contributions

U.D., V.R.Y., and A.K. conceived and designed the study. U.D. performed all computational analyses, implemented the modelling framework, and drafted the manuscript. G.S.M. contributed to data analysis. R.K., V.R.Y. and A.K. contributed to data analysis and interpretation, and assisted in manuscript preparation. A.K. supervised the project. All authors reviewed, edited, and approved the final manuscript.

## Acknowledgements

The authors acknowledge institutional support from the Department of Molecular Biology and Biotechnology, Tezpur University and the Department of Biotechnology, Koneru Lakshmaiah Education Foundation. We thank the members of the Nucleix Lab for helpful discussions.

