## Supplementary file for "Quantifying the contribution of DNA conformational flexibility to transcription factor binding on nucleosomal DNA uncovers indirect readout across diverse TF families"

### Supplementary Figure Legends

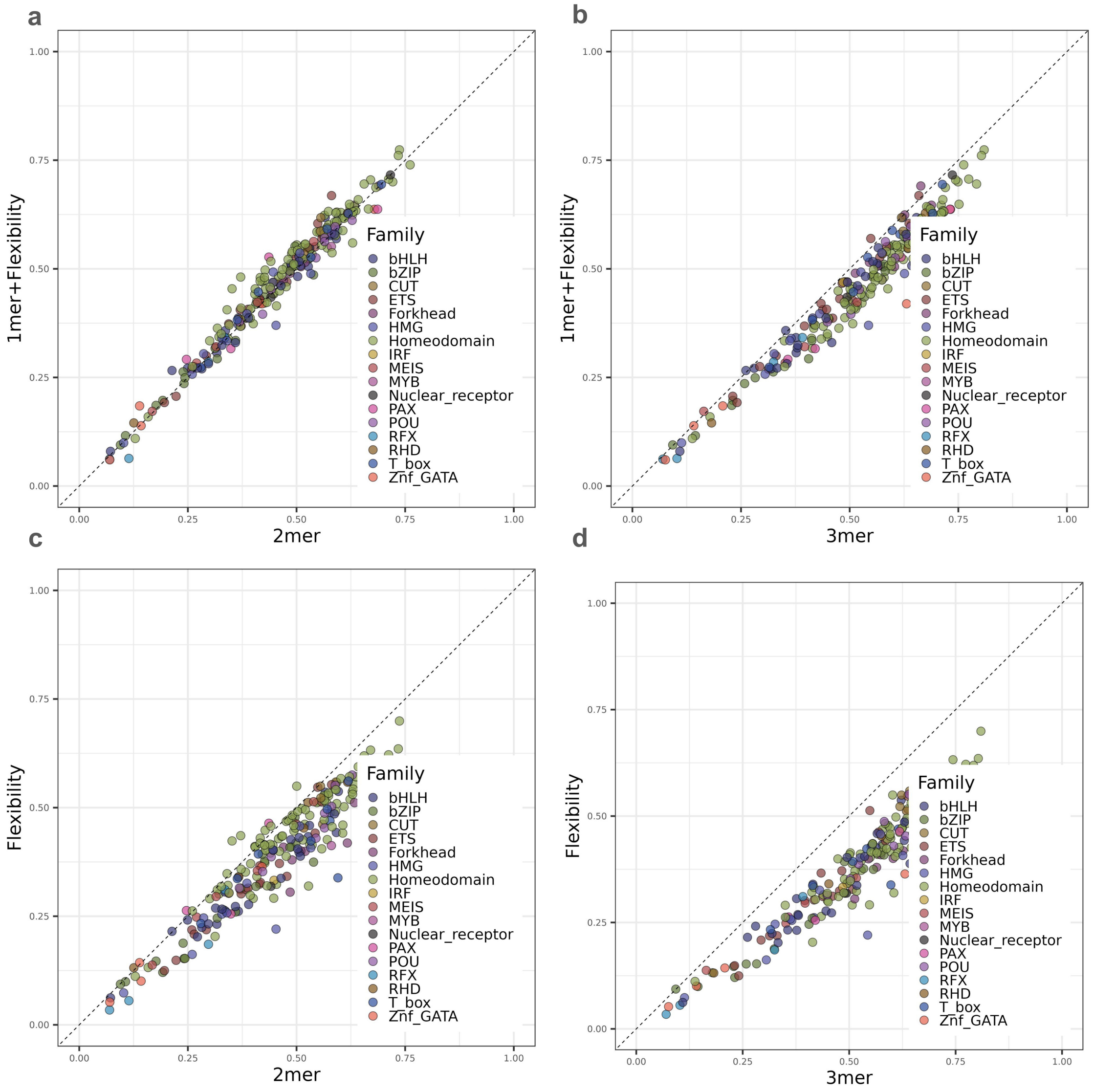

a
b
c
d
1mer+Flexibility
Flexibility
2mer
3mer
0.00
0.25
0.50
0.75
1.00
Family
bHLH
bZIP
CUT
ETS
Forkhead
HMG
Homeodomain
IRF
MEIS
MYB
Nuclear_receptor
PAX
POU
RFX
RHD
T_box
Znf_GATA


Figure 1

**Supplementary Figure S1. Higher-order *k*-mer baselines partially capture the sequence information represented by DNA flexibility descriptors in NCAP-SELEX.** (*a*) Per-dataset cross-validated $R^2$ for 2-mer models versus 1-mer + flexibility models, colored by DNA-binding domain (DBD) family. The 1-mer + flexibility model performs comparably to the larger 2-mer sequence baseline. (*b*) Equivalent comparison for 3-mer models versus 1-mer + flexibility models. The 3-mer baseline performs better overall, consistent with its richer encoding of local nucleotide dependencies. (*c*) Per-dataset comparison of 2-mer models versus flexibility-only models. (*d*) Per-dataset comparison of 3-mer models versus flexibility-only models.

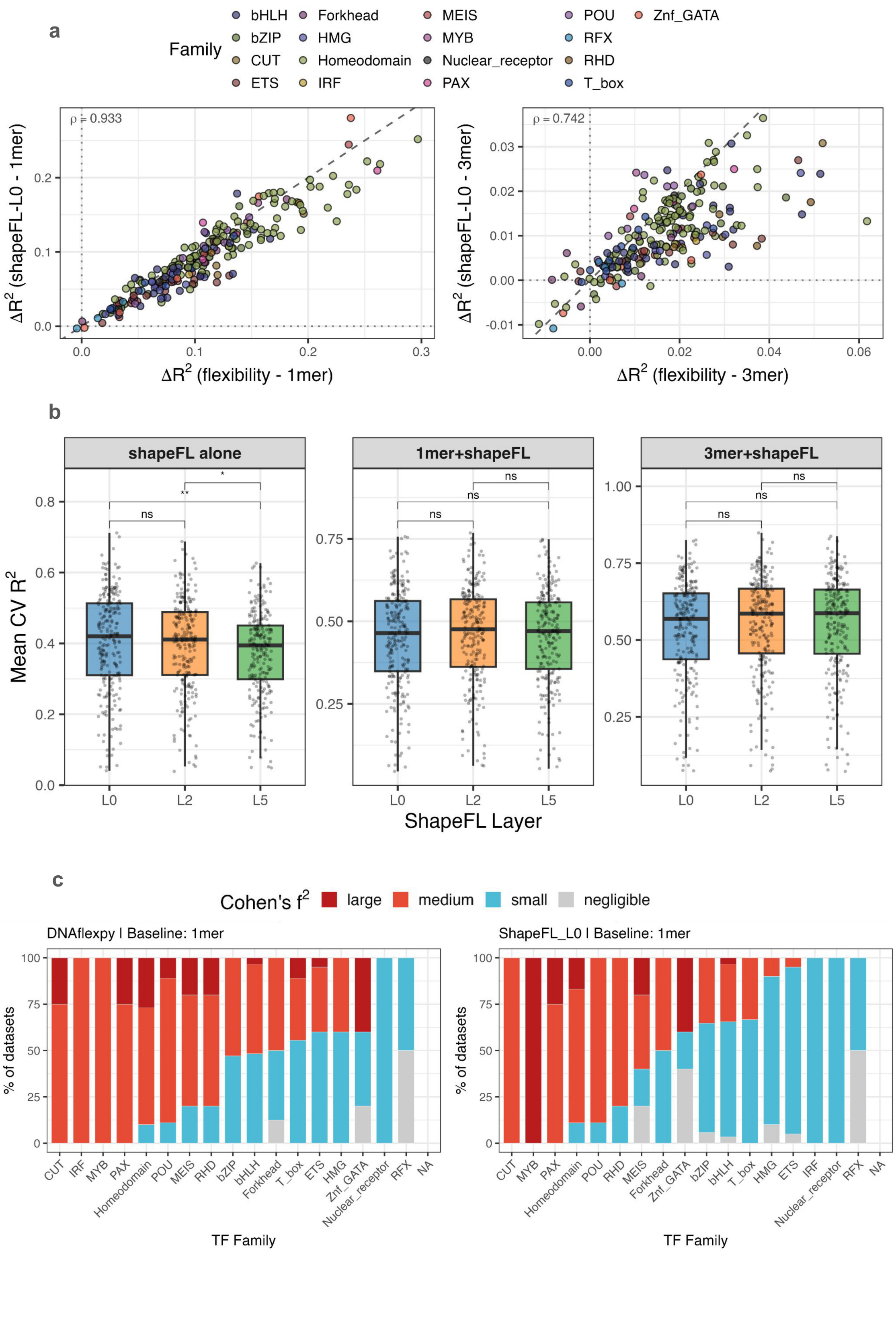

a
Family
bHLH
bZIP
CUT
ETS
Forkhead
HMG
Homeodomain
IRF
MEIS
MYB
Nuclear_receptor
PAX
POU
RFX
RHD
T_box
Znf_GATA
ρ = 0.933
ΔR² (shapeFL-L0 - 1mer)
ΔR² (flexibility - 1mer)
ρ = 0.742
ΔR² (shapeFL-L0 - 3mer)
ΔR² (flexibility - 3mer)
b
shapeFL alone
1mer+shapeFL
3mer+shapeFL
Mean CV R²
ns
**
*
ShapeFL Layer
L0
L2
L5
c
Cohen's f²
large
medium
small
negligible
DNAflexpy | Baseline: 1mer
ShapeFL_L0 | Baseline: 1mer
% of datasets
TF Family


Figure 2

**Supplementary Figure S2. DNAflexpy and Deep DNAshape shape-fluctuation features recover largely overlapping structural signal across NCAP-SELEX.** (*a*) Per-dataset $\Delta R^2$ when DNAflexpy or ShapeFL-L0 is added to the 1-mer baseline (left; Spearman $\rho = 0.933$) or the 3-mer baseline (right; Spearman $\rho = 0.742$). The two encodings identify largely similar datasets in which structural information improves prediction. (*b*) Mean cross-validated $R^2$ for ShapeFL-only, 1-mer + ShapeFL, and 3-mer + ShapeFL models across Deep DNAshape layers L0, L2, and L5, corresponding to 0-, 2-, and 5-nt extended sequence context. Increasing ShapeFL context depth adds little under this regression framework. (*c*) Per-family fraction of TF datasets in Cohen's $f^2$ categories for DNAflexpy and ShapeFL-L0 added to the 1-mer baseline. Family-level effect-size distributions are broadly concordant between the two structural representations.

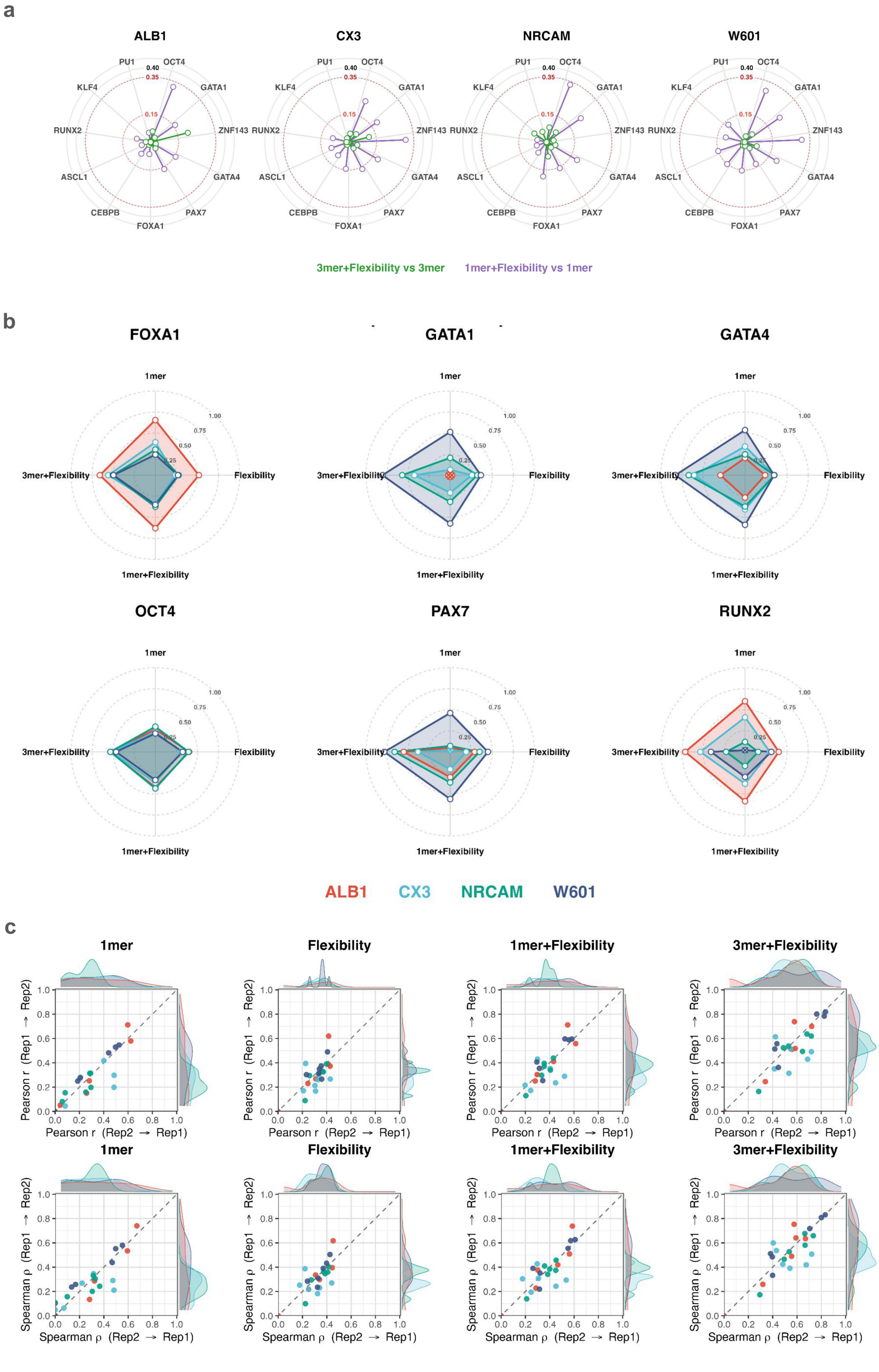
a
ALB1
CX3
NRCAM
W601
PU1
OCT4
GATA1
ZNF143
GATA4
PAX7
FOXA1
CEBPB
ASCL1
RUNX2
KLF4
0.40
0.35
0.15
3mer+Flexibility vs 3mer
1mer+Flexibility vs 1mer
b
FOXA1
GATA1
GATA4
OCT4
PAX7
RUNX2
1mer
Flexibility
1mer+Flexibility
3mer+Flexibility
1.00
0.75
0.50
0.25
ALB1
CX3
NRCAM
W601
c
1mer
Flexibility
1mer+Flexibility
3mer+Flexibility
Pearson r (Rep1 → Rep2)
Pearson r (Rep2 → Rep1)
Spearman ρ (Rep1 → Rep2)
Spearman ρ (Rep2 → Rep1)

Figure 3

**Supplementary Figure S3. Performance for $L_2$-regularized models on PIONEAR-seq data.** ($a$) Per-template radar plots of cross-validated $R^2$ across PIONEAR-seq TFs for ALB1, CX3, NRCAM, and W601. Each vertex represents a TF. Polygons compare 3-mer + flexibility versus 3-mer and 1-mer + flexibility versus 1-mer, highlighting template-dependent flexibility gains. ($b$) Per-TF radar plots of template-resolved performance for representative TFs: FOXA1, GATA1, GATA4, OCT4, PAX7, and RUNX2. Each vertex represents a model class, and each colored polygon represents one nucleosomal template. The 3-mer + flexibility encoding gives the most reproducible profile across templates, whereas flexibility-only and 1-mer models are more dispersed. ($c$) Cross-replicate concordance for each PIONEAR-seq model class. Scatter plots compare reciprocal train-test replicate directions for Pearson $r$ and Spearman $\rho$ under 1-mer, flexibility-only, 1-mer + flexibility, and 3-mer + flexibility models. Points are colored by template, with marginal densities shown along both axes. Replicate stability increases from simple sequence or flexibility-only encodings toward the 3-mer + flexibility model.

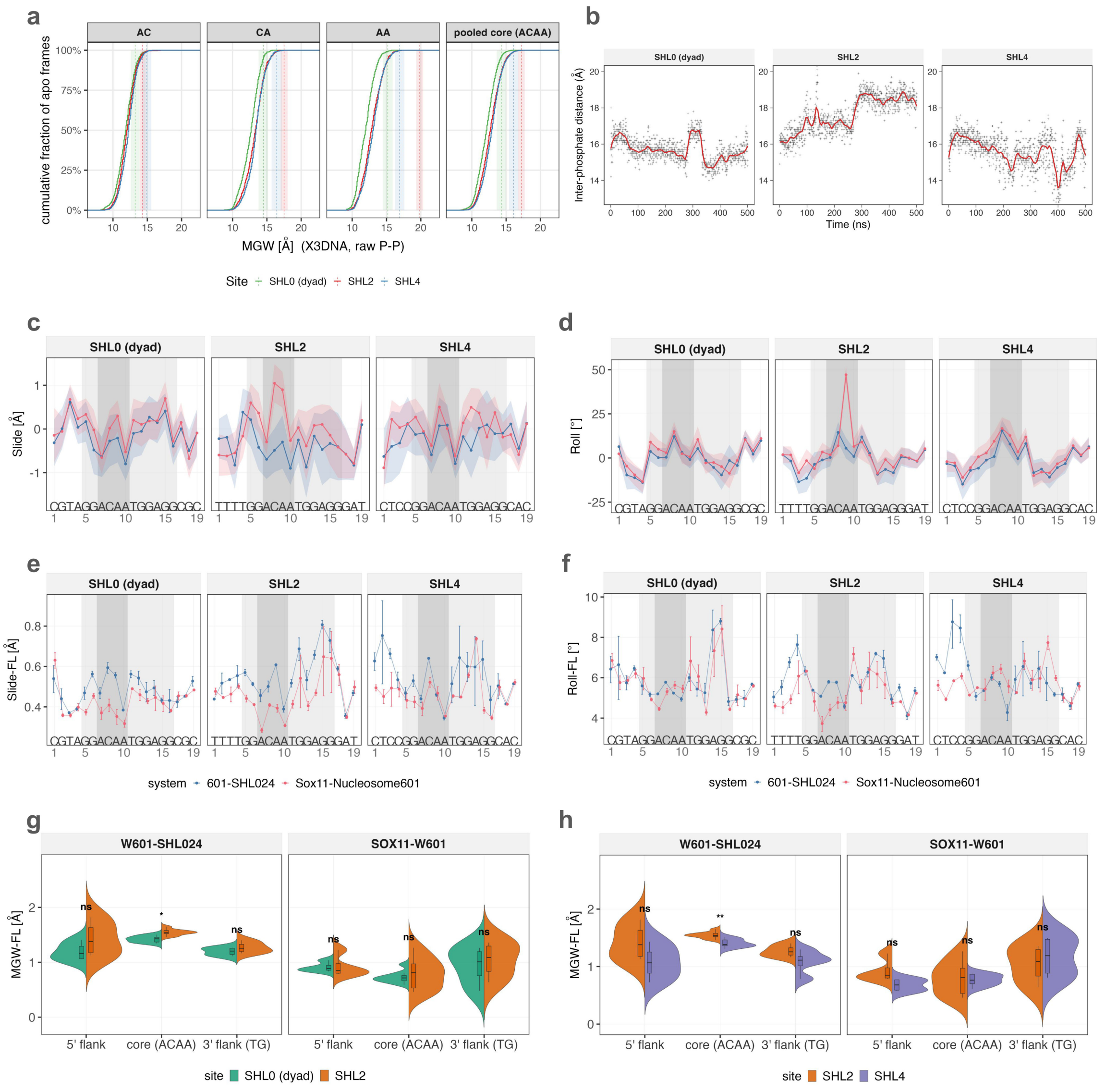

Figure 4

**Supplementary Figure S4. Atomistic features supporting SOX-nucleosome indirect readout at SHL2.** (*a*) Cumulative distributions of per-frame minor groove width (MGW; calculated from X3DNA) at the core AC, CA, AA, and pooled ACAA steps of the inserted SOX motif in SOX-unbound W601-SHL024 trajectories, stratified by SHL0, SHL2, and SHL4. The SOX11-bound geometry is indicated for reference with colored lines. Unbound trajectories rarely sample the expanded MGW geometry observed in the bound state. (*b*) Time-resolved inter-phosphate distance for CA step, at the inserted motif across SHL0, SHL2, and SHL4, shown as per-frame values with LOESS trends. (*c*) Per-position Slide profiles across the SOX motif for unbound and SOX11-bound nucleosomes at SHL0, SHL2, and SHL4. (*d*) Per-position Roll profiles for the same systems; the strongest bound-state perturbations occur near the motif core and flanks. (*e*) Per-position Slide fluctuation profiles for W601-SHL024 (SOX unbound nucleosome), and SOX11-bound nucleosomes. (*f*) Equivalent Roll fluctuation profiles for the same systems. (*g*) MGW-fluctuation distributions at the 5′ flank, central ACAA core, and 3′ flank for unbound and SOX11-bound trajectories, comparing SHL0 and SHL2. (*h*) Equivalent comparison for SHL2 and SHL4. The SHL2 core shows elevated pre-binding MGW fluctuation and the strongest post-binding reduction, consistent with SHL2 as the preferred register for SOX engagement.

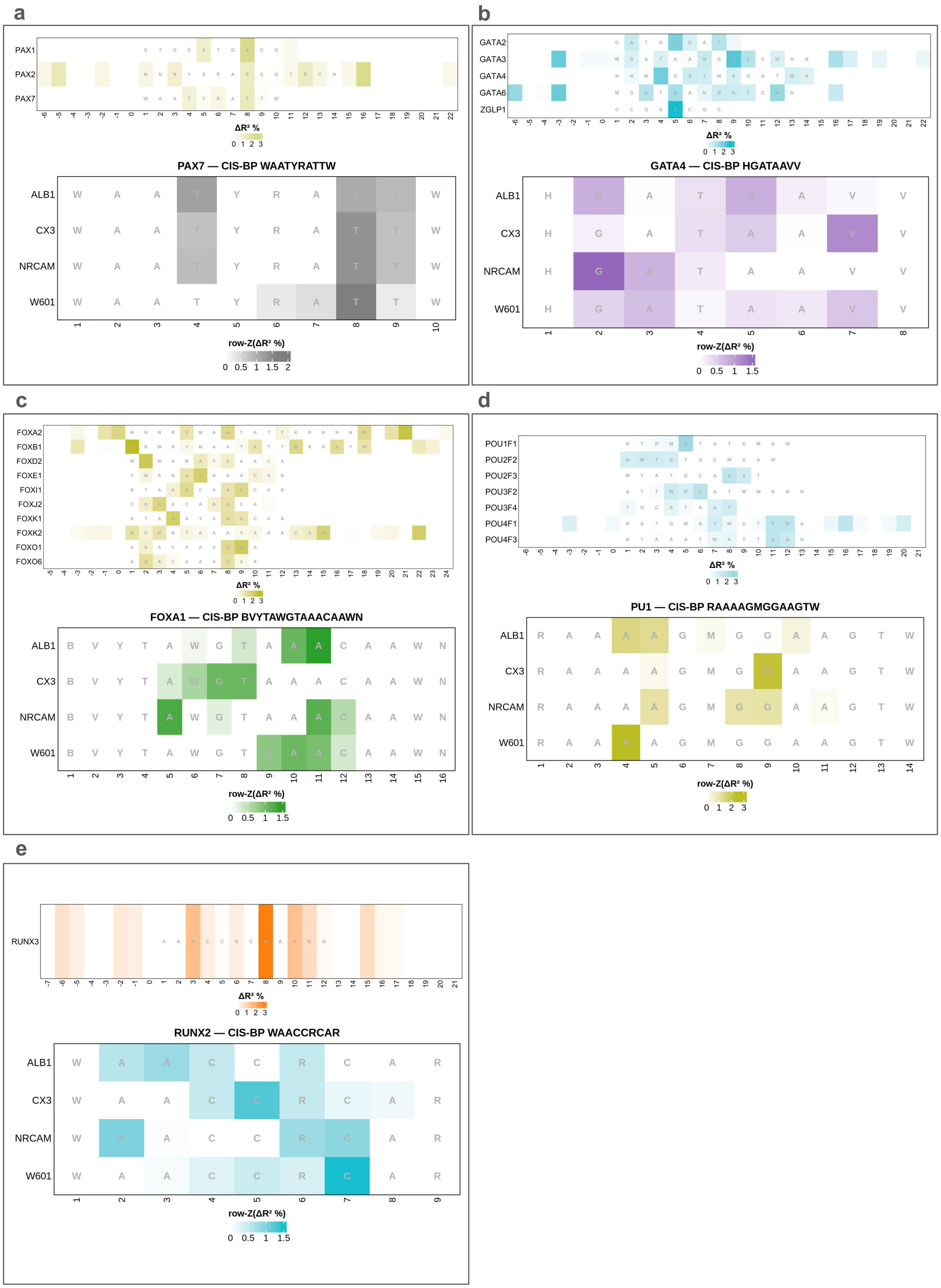
a
PAX7 — CIS-BP WAATYRATTW
b
GATA4 — CIS-BP HGATAAVV
c
FOXA1 — CIS-BP BVYTAWGTAAACAAWN
d
PU1 — CIS-BP RAAAAGMGGAAGTW
e
RUNX2 — CIS-BP WAACCRCAR
ALB1
CX3
NRCAM
W601
row-Z(ΔR² %)
ΔR² %

Figure 5

**Supplementary Figure S5. Model inferred flexibility footprints for NCAP-SELEX and PIONEAR-seq TFs.** Each panel shows two flexibility footprint profiles for a TF familiy ( *top:* model trained and evaluated with NCAP-SELEX data; *bottom:* model trained and evaluated with PIONEAR-seq data for a TF across different templates). The top heatmap shows position-resolved flexibility contribution across sequences aligned to the indicated NCAP-SELEX motif. The bottom heatmap shows template-resolved footprint structure for a TF from the same family, with rows corresponding to ALB1, CX3, NRCAM, and W601 and columns corresponding to motif positions. Color reports normalized $\Delta R^2$ gain from adding flexibility above the 1-mer baseline (Methods). (*a*) Flexibility footprints for PAX TF family factors with aligned motifs from NCAP-SELEX study (*top*); flexibility footprint for PAX7 TF across templates with CIS-BP motif WAATYRATTW (*bottom*). (*b*) Same profile for TFs from GATA family, and GATA4 with CIS-BP motif HGATAAVV. (*c*) TFs from Forkhead family, FOXA1 TF CIS-BP motif BVYTAWGTAAACAAWN. (*d*) TFs with PU domain, PU.1 with CIS-BP motif RAAAAGMGGAAGTW. (*e*) TFs from RUNX families, RUNX2 with CIS-BP motif WAACCRCAR.

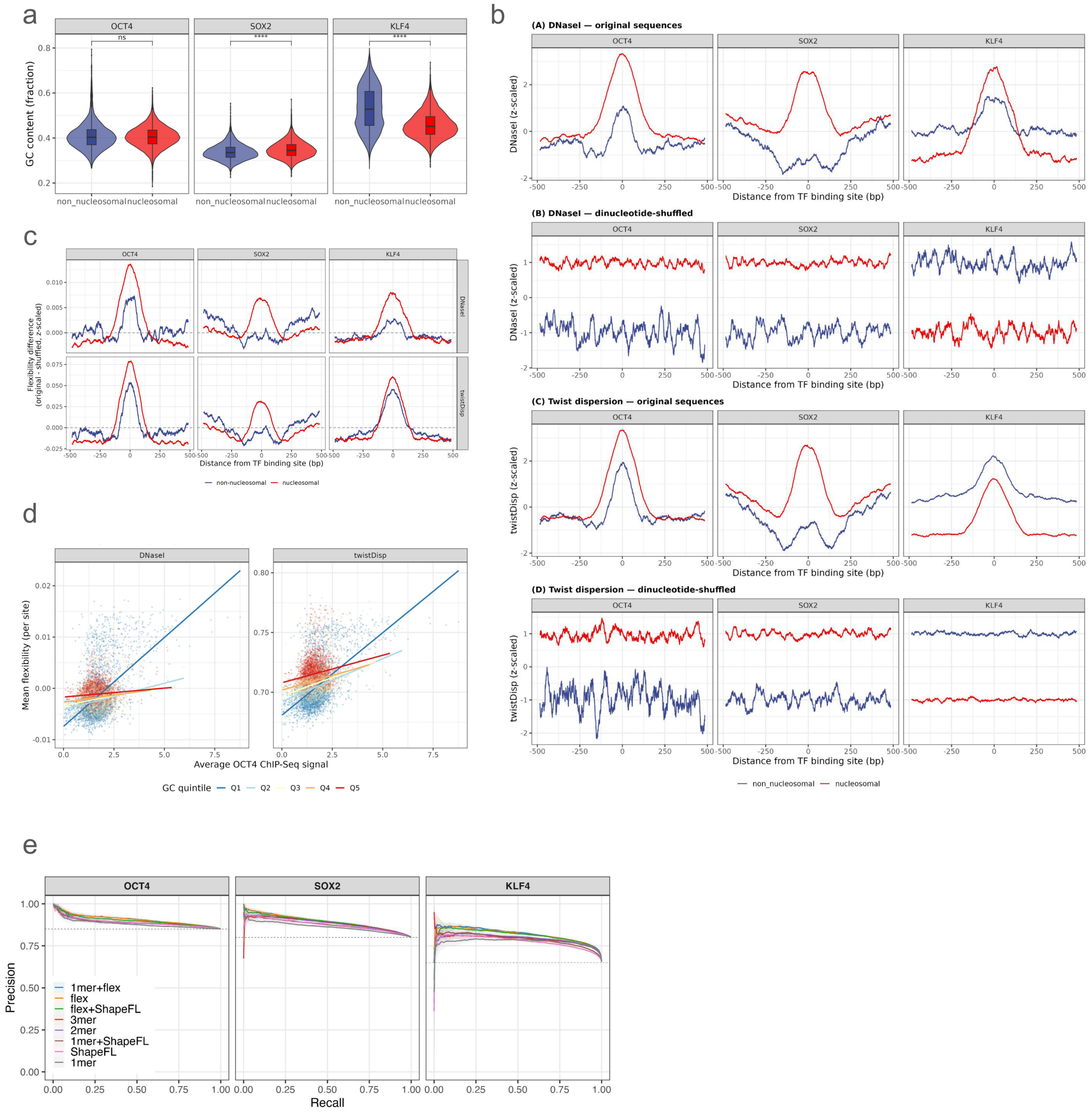
a
OCT4
SOX2
KLF4
GC content (fraction)
non_nucleosomal nucleosomal
b
(A) DNaseI — original sequences
(B) DNaseI — dinucleotide-shuffled
(C) Twist dispersion — original sequences
(D) Twist dispersion — dinucleotide-shuffled
DNaseI (z-scaled)
twistDisp (z-scaled)
Distance from TF binding site (bp)
non_nucleosomal nucleosomal
c
Flexibility difference (original - shuffled, z-scaled)
DNaseI
twistDisp
non-nucleosomal nucleosomal
d
Mean flexibility (per site)
Average OCT4 ChIP-Seq signal
GC quintile Q1 Q2 Q3 Q4 Q5
e
Precision
Recall
1mer+flex
flex
flex+ShapeFL
3mer
2mer
1mer+ShapeFL
ShapeFL
1mer

Figure 6

**Supplementary Figure S6. Control analyses supporting in vivo flexibility associations at OSK and GATA3 binding sites.** (*a*) GC-content distributions for nucleosomal and non-nucleosomal OCT4, SOX2, and KLF4 binding sites in IMR90 cells. OCT4 shows no significant GC-content difference between groups (40.5% versus 40.9%; Wilcoxon rank-sum test, $p = 0.528$; Cliff's $\delta = 0.004$), SOX2 shows a small shift (Cliff's $\delta = 0.109$), and KLF4 shows a larger GC-content bias (45.8% versus 53.1%; Cliff's $\delta = 0.380$). (*b*) Average DNase I sensitivity and twist-dispersion profiles across OCT4, SOX2, and KLF4 binding sites in nucleosomal and non-nucleosomal regions, comparing original sequences with dinucleotide-preserving shuffled controls generated using *bias-away* ($k = 2$). Binding-site-centered flexibility peaks are reduced or abolished in shuffled controls, indicating position-specific sequence organization rather than dinucleotide composition alone. (*c*) Normalized average flexibility profiles for OCT4, SOX2, and KLF4 sites stratified by nucleosomal binding sites. (*d*) Mean local DNA flexibility versus mean OCT4 ChIP-seq signal across IMR90 nucleosomal binding sites for DNase I sensitivity and twist dispersion, with points colored by GC-content quintile. Lines show linear fits. (*e*) Precision-recall curves for cross-validated XGBoost models classifying OSK binding sites as nucleosomal or non-nucleosomal, complementing the ROC analysis in Fig. 5d. Feature sets include 1-mer, 2-mer, 3-mer, flexibility-only, ShapeFL-only, 1-mer + flexibility, 1-mer + ShapeFL, and flexibility + ShapeFL.

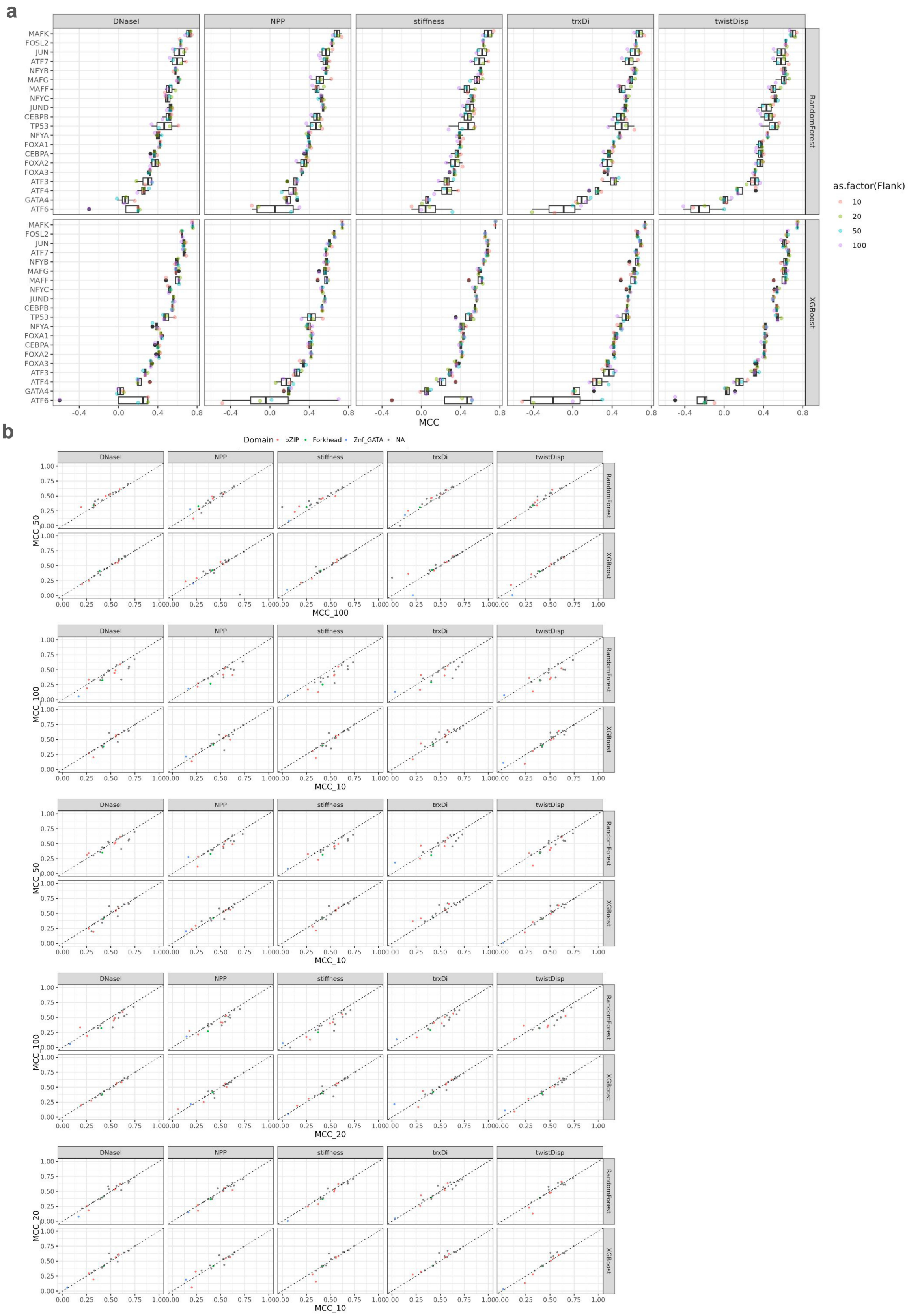
a
DNaseI
NPP
stiffness
trxDi
twistDisp
RandomForest
XGBoost
MAFK
FOSL2
JUN
ATF7
NFYB
MAFG
MAFF
NFYC
JUND
CEBPB
TP53
NFYA
FOXA1
CEBPA
FOXA2
FOXA3
ATF3
ATF4
GATA4
ATF6
MCC
as.factor(Flank)
10
20
50
100
b
Domain
bZIP
Forkhead
Znf_GATA
NA
MCC_50
MCC_100
MCC_10
MCC_20

Figure 7

**Supplementary Figure S7. Extended ENCODE classification results across PTFs, flexibility descriptors, classifiers, and motif-flank sizes.** (*a*) Matthews correlation coefficient (MCC) per TF for each DNAflexpy descriptor added to the 1-mer sequence baseline. Results are shown for Random Forest and XGBoost classifiers. Point color indicates the motif-flank size used for feature extraction: 10, 20, 50, or 100 bp. (*b*) MCC scatter matrix comparing classification performance across motif-flank sizes. Each column corresponds to one flexibility descriptor, and each row block corresponds to one classifier. Within each panel, the two axes compare MCC values obtained at different flank sizes, with points colored by DBD family. Concentration along the diagonal indicates that model performance is relatively stable across flank-size choices; torsional and rotational descriptors, especially twist dispersion and trx, show the most stable transfer across these settings.
